# Nucleotide inhibition of the pancreatic ATP-sensitive K+ channel explored with patch-clamp fluorometry

**DOI:** 10.1101/803999

**Authors:** Samuel G. Usher, Frances M. Ashcroft, Michael C. Puljung

**Affiliations:** Department of Physiology, Anatomy and Genetics, University of Oxford, Oxford OX1 3PT, United Kingdom

## Abstract

Pancreatic ATP-sensitive K+ channels (K_ATP_) comprise four inward rectifier subunits (Kir6.2), each associated with a sulphonylurea receptor (SUR1). ATP/ADP binding to Kir6.2 shuts K_ATP_. Mg-nucleotide binding to SUR1 stimulates K_ATP_. In the absence of Mg^2+^, SUR1 increases the apparent affinity for nucleotide inhibition at Kir6.2 by an unknown mechanism. We simultaneously measured channel currents and nucleotide binding to Kir6.2. Fits to combined data sets suggest that K_ATP_ closes with only one nucleotide molecule bound. A Kir6.2 mutation (C166S) that increases channel activity did not affect nucleotide binding, but greatly perturbed the ability of bound nucleotide to inhibit K_ATP_. Mutations at position K205 in SUR1 affected both nucleotide affinity and the ability of bound nucleotide to inhibit K_ATP_. This suggests a dual role for SUR1 in K_ATP_ inhibition, both in directly contributing to nucleotide binding and in stabilising the nucleotide-bound closed state.

## Introduction

ATP-sensitive K+channels (K_ATP_) couple the metabolic state ofacellto its electrical activity (Ashcroft and Rorsman, 2013). In pancreatic β-cells, closure of K_ATP_ in response to glucose uptake triggers insulin secretion. As such, mutations in K_ATP_ that affect its response to changes in cellular metabolism cause diseases of insulin secretion, e.g. neonatal diabetes and persistent hyperinsulinémie hypoglycaemia of infancy (PHHI; Quan et al. (2011); Ashcroft et al (2017)). K_ATP_ is composed of four inwardly rectifying K+ channel subunits (Kir6.2 in pancreatic β-cells), which form the channel pore and four modulatory sulphonyiurea receptor subunits (SUR1 in β-cells; Figure 1A; Aguilar-Bryan et al. (1995); Inagaki et al. (1995); Sakura et al. (1995); Inagaki et al (1997)). SUR1 is a member of the ABC transporter family but lacks any transport activity (Aguilar-Bryan et al. 1995; Tusnady et al., 1997). K_ATP_ responds to metabolism via adenine nucleotide binding to three distinct classes of intracellular nucleotide-binding site (one on each Kir6.2 subunit and two on each SUR1 subunit—making twelve sites in total (Vedovato et al., 2015). Binding of ATPor ADP to Kir6.2 inhibits K_ATP_ channel activity (Tucker et al., 1997; Proks et al., 2010), whereas binding of nucleotides to SUR1 stimulates K_ATP_ (Nichols et al., 1996; Tucker et al., 1997)). The stimulatory activity of nucleotides on K_ATP_ depends on Mg^2+^ (Gribble et al., 1998), whereas their inhibitory effect on Kir6.2 does not (Tucker et al., 1997).

**Figure 1.**
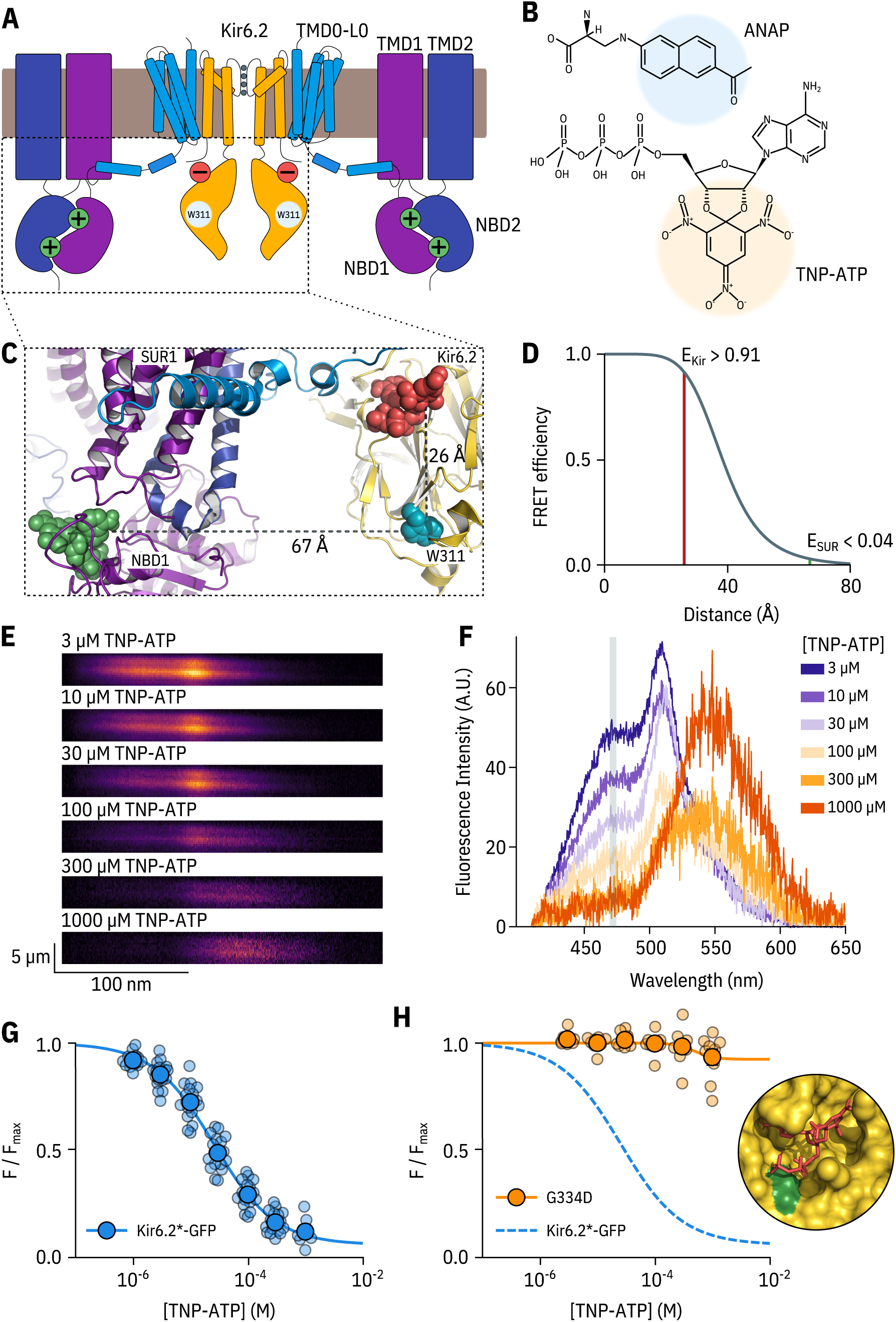
A FRET assay to measure nucleotide binding to Kir6.2. **A.** Cartoon illustrating the topology of K_ATP_, The inhibitory nucleotide-binding site on Kir6.2 is shown in red; the stimulatory nucleotide-binding sites on SUR1 are shown in green. The three transmembrane domains of SUR1 are designated TMDO, TMD1, and TMD2. The loop connecting TMDO to TMD1 is designated LO. The nucleotide binding domains of SUR1 are labelled NBD1 and NBD2. **B.** Chemical structures of ANAP and TNP-ATP. The fluorescent moieties are highlighted. **C.** Side view of the structure of the cytosolic domains of Kir6.2 (PDB accession #6BAA)and one SUR1 subunit (PDB accession #6PZI). TNP-ATP (red, from PDB accession #5XW6) was docked into the nucleotide-binding site of Kir6.2 and positioned in NBS1 of SUR1 (green, from PDB accession #3AR7) by alignment as described in Materials and Methods. Distances from the native tryptophan at position 311 in Kir6.2 to the fluorescent moieties of the TNP-ATPs are displayed in A. D. Theoretical FRET efficiency between ANAP and TNP-ATP as a function of distance, calculated from the Forster equation. The distances and corresponding FRET efficiencies between ANAP at position 311 and TNP-ATP bound to Kir6.2 (E_K_j_r_) and SUR1 (E_SUR_) are indicated. **E.** Spectral images acquired from an unroofed membrane expressing Kir6.2*-GFP + SUR1 and exposed to increasing concentrations of TNP-ATP. The y-dimension in each image represents distance. The x-dimension represents wavelength. **F.** Line-averaged, background-subtracted spectra from **E** displayed with increasing concentrations of TNP-ATP coloured from purple to orange. The three fluorophores have distinct peaks: ANAP at 472 nm, GFP at 508 nm, and TNP-ATP at 561 nm. The shaded rectangle indicates the wavelength range used to measure ANAP intensity. **G.** Concentration-response relationship for binding of TNP-ATP to Kir6.2*-GFP + SUR1 in unroofed membranes. Data were plotted as *F*/*F_max_*, where *F_max_* is the fluorescence intensity in the absence of nucleotide. The smooth curve is a descriptive Hill fit. *EC*_50_ *=* 25.6 μΜ, *h =* 0.82, *E_max_ =* 0.93, n = 18. **H.** Concentration-response relationship for binding of TNP-ATP to Kir6.2*-G334D-GFP + SUR1 in unroofed membranes. The dashed blue curve is the fit from G. The orange curve is a descriptive Hill fit to the G334D data. *EC*_50_ *=* 493 μΜ, *h =* 2.63, *E_max_ =* 0.08, n = 9. The inset shows the location of G334D (green) in relation to the inhibitory ATP binding site on Kir6.2 (PDB accession #6BAA). TNP-ATP (PDB accession #5XW6) shown in red sticks.

In addition to nucleotide-dependent activation, SUR1 confers several other properties on the Kir6.2. First, association with SUR1 increases the open probability (*P_open_*) of Kir6.2 (Babenko and Bryan, 2003; Chan et al., 2003; Fang et al., 2006). Despite this increase in *P_open_*, SUR1 also paradoxically increases the apparent affinity for nucleotide inhibition at Kir6„2 by an unknown mechanism (Tucker et al., 1997). SUR1 is also responsible for high-affinity inhibition of K_ATP_ by antidiabetic sulphonyl ureas and glinidesas well as activation by K_ATP_-specific K^+^ channel openers (Tucker et al., 1997). Finally, SUR1 and Kir6.2 must co-assemble to ensure mutual exit from the endoplasmic reticulum and correct trafficking to the plasma membrane (Zerangue et al., 1999).

To date, the primary means of studying nucleotide-dependent effects on K_ATP_ channel function has been with electrophysiological approaches, which measure the summed activity of all three classes of binding site acting in concert. Thus, it can be difficult to separate the contributions of each class of site to the opening and closing of the channel pore and to properly distinguish between nucleotide binding and channel gating. To overcome these limitations, we have applied a novel approach to directly measure nucleotide binding to each individual class of site in K_ATP_ (Puljung et al., 2019). This method utilizes Forster resonance energy transfer (FRET) between channels labelled with the fluorescent unnatural amino acid 3-(6-acetylnaphthalen-2-ylamino)-2-aminopropanoic acid (ANAP) and fluorescent tri nitrophenyl (TNP) analogues of adenine nucleotides (Figure 1B). As we show here, this method is readily combined with patch-clamp electrophysiology so that nucleotide binding and regulation of current can be measured simultaneously. This has enabled us to quantitatively assess nucleotide binding to Kir6.2 and explore how this is coupled to channel inhibition in both wild-type K_ATP_ and K_ATP_ carrying mutations that impair ATP inhibition.

## Results

### Measuring nucleotide binding to Kir6.2

We previously used this FRET-based binding assay to measure nucleotide binding to the second nucleotide-binding site of SUR1 (Puljung et al., 2019). To measure binding to Kir6.2 in the complete K_ATP_ complex (four full-length Kir6.2 subunits co-expressed with four full-length SUR1 subunits), we replaced a tryptophan at position 311 (W311) that is 26 Λ from the location of the inhibitory nucleotide-binding site on Kir6.2 with ANAP (Figure 1C) such that each subunit is labelled with one ANAP molecule. We designate this construct Kir6.2*. Based on the theoretical FRET efficiency calculated from the Forster equation and available cryo-EM structures (Martin et al., 2017,2019)*, we expect* 91 % FRET efficiency between ANAP at position 311 and a TNP-ATP molecule bound to Kir6.2, and only 4% FRET efficiency to TNP-ATP bound to the closest nucleotide-binding site on SUR1 (nucleotide binding site 1, Figure 1D). We also expect very little FRET between ANAP at position 311 and TNP-ATP bound to neighbouring Kir6.2 subunits (15-25%).

ANAP incorporation into Kir6.2 was achieved as described previously (Chatterjee et al., 2013; Zagotta et al., 2016; Puljung et al., 2019). Briefly, HEK-293T cells were co-transfected with a plasmid encoding a Kir6.2 construct with a C-terminal GFP tag and an amber stop codon (TAG) replacing the codon corresponding to amino acid position 311 (W31 1^tag^-GFP) and a plasmid encoding an ANAP-specific tRNA/tRNA synthetase pair (pANAP). We also included a dominant negative eukaryotic ribosomal release factor (eRF-E55D) in our transfections, which has been shown to increase the amount of full-length, ANAP-labelled protein (Schmied et al., 2014; Puljung et al., 2019). When cultured in the presence of ANAP, full length, fully ANAP-labelled Kir6.2 protein was produced and successfully trafficked to the membrane in the presence of SUR 1 (Figure 1—Figure supplement 1; see Materials and Methods). ANAP fluorescence from labelled channels can be separated from unincorporated ANAP or autofluorescence based on emission spectra (Puljung et al., 2019). However, we found it much more convenient to first identify transfected cells or membrane fragments based on the presence of a GFP tag. Thus, we used GFP-tagged Kir6.2 constructs throughout this study, unless otherwise indicated.

In all our experiments, we measured currents in excised patches from cells expressing K_ATP_ in the absence of Mg^2+^. Under such conditions, nucleotides can bind to both sites on SUR1, but no activation occurs, allowing inhibitory currents to be measured in isolation (Gribble et al., 1998; Ueda et al., 1999; Puljung et al., 2019). Kir6.2*-GFP + SUR1 exhibited nearly identical sensitivity to ATP inhibition as Kir6.2-GFP + SUR1 (Figure 1—Figure supplement 2A), indicating that replacement of W311 with ANAP did not affect inhibition by K_ATP_. Both subunits also showed similar sensitivity to TNP-ATP, which inhibited with a higher apparent affinity relative to ATP (Figure 1—Figure supplement 2B,C).

Kir6.2-GFP has been demonstrated to traffic to the plasma membrane in the absence of SUR1 (John et al., 1998; Makhina and Nichols, 1998). In a luminescence-based, surface-expression assay, we did not detect HA-tagged Kir6.2*-GFP at the plasma membrane in the absence of SUR1 (Figure 1—Figure supplement 1E). To verify that the currents measured in our experiments in which Kir6.2*-GFP was co-transfected with SUR1 were the result of Kir6.2*-GFP + SUR1 and not Kir6.2*-GFP alone, we measured the sensitivity of currents to inhibition by the sulphonyiurea tolbutamide, a property conferred by the SUR1 subunit. Whereas currents from unlabelled wild-type Kir6.2-GFP expressed in the absence of SUR1 were not affected by 100 μΜ tolbutamide, both wild-type Kir6.2-GFP and Kir6.2*-GFP currents were inhibited to a similar extent by when expressed with SUR1 (46.5% ±0.04% and 57.7% ±0.02%, respectively; Figure 1—Figure supplement 2D). The extent of block was similar to previous measurements of tolbutamide inhibition (Tucker et al., 1997), confirming that Kir6.2*-GFP was co-assembled with SUR1 at the plasma membrane.

To measure nucleotide binding, cells transfected with Kir6.2*-GFP + SUR1 were briefly sonicated, leaving behind unroofed plasma membrane fragments (Heuser, 2000; Zagotta et al, 2016; Puljung et al., 2019) contain ingANAP-labelled K_ATP_ channels with the intracellular nucleotide-binding sites exposed to the bath solution. The sample was excited with a 385 nm LED and emitted fluorescence from the membrane fragments was passed through a spectrometer, allowing us to separate ANAP, GFP, and TNP-ATP fluorescence by peak wavelength (Figure 1 E,F). As expected from FRET, increasing the concentration of TNP-ATP caused a decrement in the ANAP peak at 472 nm and a concomitant increase in the TNP-ATP peak at 561 nm (Figure 1F). We used the quenching of the ANAP peak as a direct measure of TNP-ATP binding as this signal was specific to K_ATP_. In contrast, the peak TNP-ATP fluorescence may include contributions from both specific and non-specific nucleotide binding. Due to the sharp cut-off of the GFP emission spectrum at shorter wavelengths, our measurements of peak ANAP fluorescence were unaffected by the presence of the GFP tag on Kir6.2.

We fit concentration-response data for TNP-ATP quenching with the Hill equation, to produce estimates of apparent affinity and E_max_ (ANAP quenching at saturating concentrations of TNP-ATP; Figure 1G). E_max_ was 93%, in good agreement with the 91% predicted by the Forster equation and theoretical distance measurements (Figure 1D), suggesting that we were able to measure binding directly to the inhibitory site at Kir6.2. To confirm this, we introduced a well-studied neonatal diabetes mutation (G334D) into the Kir6.2 binding site, which drastically reduces the sensitivity of the channel to inhibition by nucleotides (Drain et al., 1998; Masia et al., 2007; Proks et al., 2010). Based on the cryo-electron microscopy structures of K_ATP_, this mutation is expected to interfere with nucleotide binding directly (Figure 1H inset, Martin et al., (2017)). The resulting construct Kir6.2*,G334D-GFP + SUR1 displayed drastically reduced ANAP quenching over the range of TNP-ATP concentrations tested, with an E_max_ of 8%; again in good agreement with the predicted FRET efficiencies between ANAP at position 311 and TNP-ATP bound only toSURI. We therefore conclude that our binding measurements were specific for the inhibitory nucleotide-binding site on Kir6.2. This observation is consistent with the interpretation that the G334D mutation causes neonatal diabetes by preventing nucleotide binding. However, the observed loss in nucleotide-dependent quenching in Kir6.2*-G334D-GFP may also be due to an allosteric effect of the G334D mutation on channel gating. We feel that this interpretation is unlikely, as G334D has been shown to have no effect on the unliganded *P_open_* of K_ATP_ (Proks et al., 2010).

### Measuring current inhibition and nucleotide binding simultaneously

The apparent affinity of Kir6.2*-GFP + SUR1 for TNP-ATP in unroofed membranes was 25.6 μΜ (Figure 1G and Table 1). This value is higher than the apparent affinity for nucleotide inhibition (6.2 μΜ) measured using patch-ciamp (Figure 1—Figure supplement 2C). However, both binding and current measurement are a function of the intrinsic binding affinity, the channel *P_open_*, and the ability of agonist, once bound, to close the channel. Furthermore, the functional state of K_ATP_ in unroofed membranes is unclear. This is a particular problem with K_ATP_ channels, which run down due to slow dissociation of phosphatidylinositol 4,5-bisphosphate (PIP_2_), reducing the *P_open_* over time even in the absence of nucleotides (Proks et al., 2016).

**Table 1.** Hill fit parameters from unroofed membranes. EC_50_ values and their standard errors are reported in log_10_ *M*^−1^.

**Tables and supplementary figures**
| Fluorescence Quenching | Construct | Term | Estimate | Standard Error |
| --- | --- | --- | --- | --- |
| *TNP-ATP* | Kir6.2*-GFP+SUR1<br>n = 18 | $EC_{50}$ | -4.59 | 0.05 |
| | | $h$ | 0.82 | 0.05 |
| | | $E_{max}$ | 0.93 | 0.03 |
| | Kir6.2*-G334D-GFP+SUR1<br>n = 9 | $EC_{50}$ | -3.31 | 2.23 |
| | | $h$ | 2.63 | 17.70 |
| | | $E_{max}$ | 0.08 | 0.26 |
| | Kir6.2*-C166S-GFP+SUR1<br>n = 12 | $EC_{50}$ | -4.50 | 0.05 |
| | | $h$ | 0.92 | 0.08 |
| | | $E_{max}$ | 0.87 | 0.03 |
| | Kir6.2*-GFP<br>n = 14 | $EC_{50}$ | -4.42 | 0.05 |
| | | $h$ | 0.83 | 0.05 |
| | | $E_{max}$ | 0.92 | 0.03 |

As measuring either nucleotide binding or ionic currents in isolation only offers limited mechanistic insight into inhibition of K_ATP_, we turned to patch-ciamp fluorometry (PCF, Proks et al., (2016); Zheng and Zagotta (2003)). Using PCF, we can measure TNP-ATP binding to Kir6.2 and channel activity simultaneously (Figure 2), providing us with direct access to the relationship between nucleotide binding and channel function. We simultaneously measured fluorescence emission spectra and ionic currents for Kir6.2*-GFP + SUR1 in inside-out, excised membrane patches. As before, all measurements were performed in the presence of Mg^2+^ chelators, such that nucleotide inhibition could be measured in the absence of activation (Tucker et al., 1997; Gribble et al., 1998). Strikingly, current inhibition occurred at a lower range of concentrations compared to nucleotide binding (Figure 2C,D). The apparent *EC*_50_ for inhibition calculated from Hill fits was an order of magnitude lower than the *EC*_50_ for binding measured in the same patches (Figure 2C, Table 2). We considered several different gating models to explain this observation. In each model, we assumed the channel pore was able to open and close in the absence of ligand with an equilibrium constant *L*, where *P_open_ = L/*(*L* + 1) and *L* > 0. This reflects the ability of K_ATP_ to open and close in the absence of nucleotides. Each model also had parameters representing the intrinsic binding affinity to the closed state (*K_A_*, where *K_A_* > 0) and the factor by which nucleotide binding favours channel closure (*D*, where *D* < 1).

**Figure 2.**
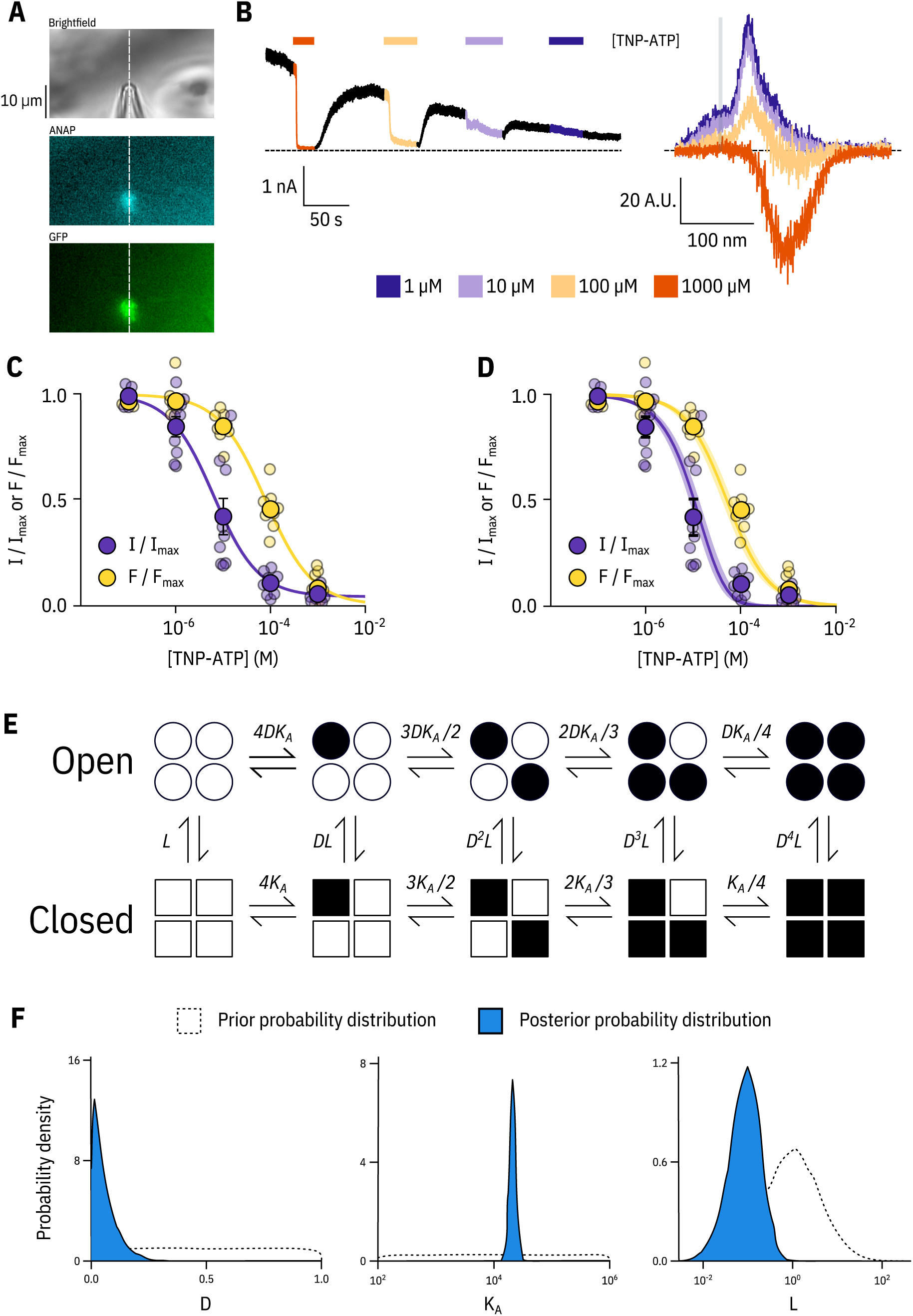
Simultaneous measurements of nucleotide binding and channel current. **A.** Brightfieid and fluorescence images of a patch pipette and excised, inside out patch expressing Kir6.2*-GFP + SUR1, with the location of the centre of the spectrometer slit overlaid as a white, vertical line. **B.** Current (left) and spectra (right) acquired from the same excised, inside-out patch exposed to TNP-ATP and coloured according to concentration. **C.** Concentration-response (n = 9) for TNP-ATP inhibition of Kir6.2*-GFP + SUR1 currents (*I*/*I_max_*) and for quenching of ANAP fluorescence (*F*/*F_max_*). Both current inhibition and fluorescence quenching were fit to the Hill equation. Current inhibition: *IC*_50_ *=* 6.23μΜ, *h =* 0.92, *I_max_*= 0.96, fluorescence quenching: *EC*_50_ = 77.7 μΜ, *h =* 0.87, *E_max_ =* 1.00. **D.** The same data as in **C** fit to an MWC-type model. Solid curves represent the median fit; shaded areas represent the 95% quantile interval. Values for the fits are reported in the text and in Table 3. **E.** MWC-type model for inhibition of K_ATP_ by nucleotides. Open subunits are shown as circles; closed are shown as squares. Nucleotide-bound subunits are represented by filled symbols. *L, D*, and *K_A_* are defined in the text. **F.** Posterior probability distributions for the MWC-type model generated by MCMC fits to the data in **C** overlaid on the prior probability distribution (dashed line) for each parameter.

**Table 2.**
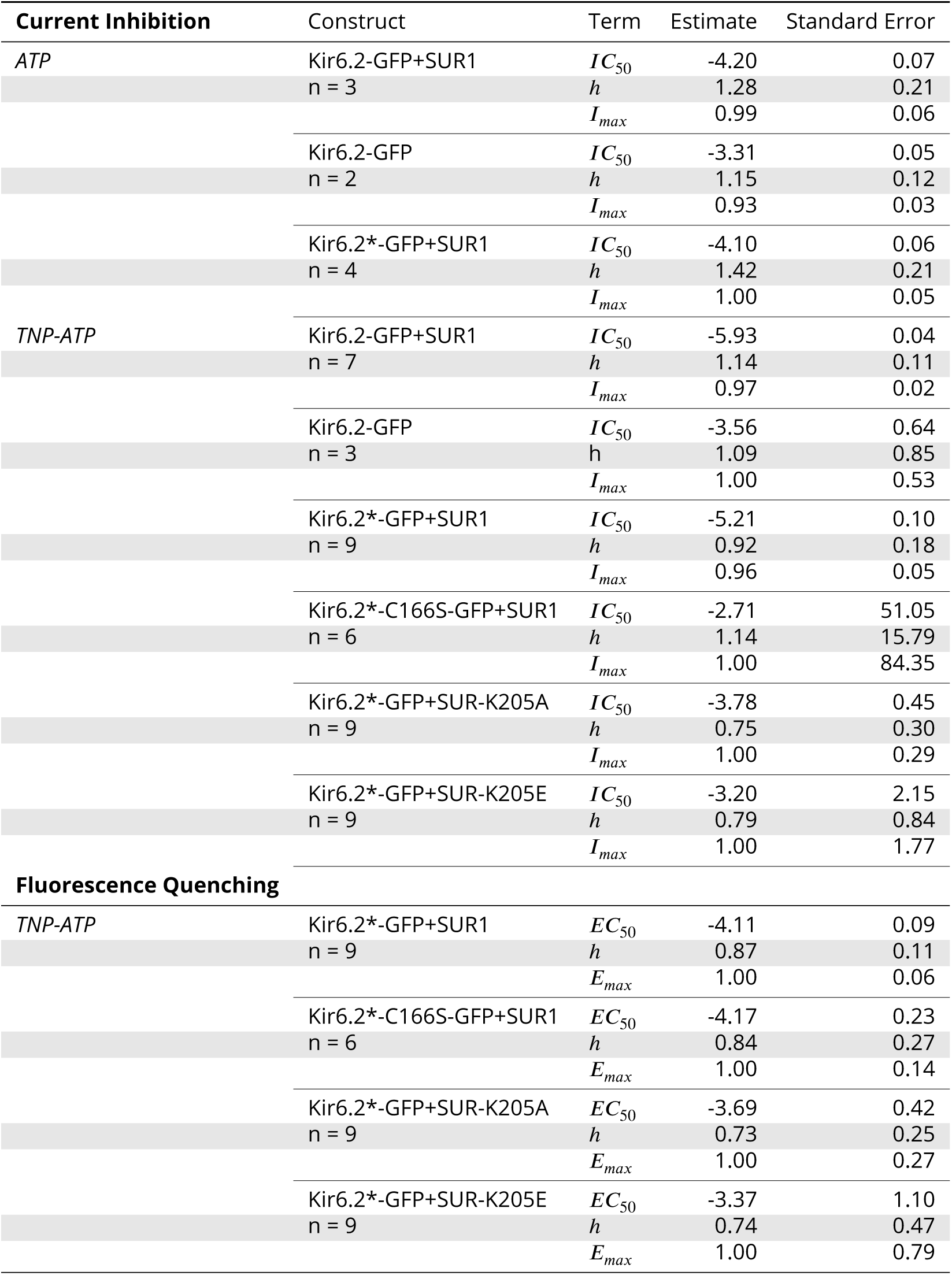
Hill fit parameters from excised patches. EC_50_ values and their standard errors are reported in log_10_ *M*^−1^.

**Table 3.** **Fitted parameters for the MWC-type models.** *L, K_A_* and their associated quantiles are reported as log_10_ values to maintain consistency of the accuracy they are reported at.

| **Full MWC** |  |  |  |  |
| --- | --- | --- | --- | --- |
| Construct | Term | Estimate | 2.5% Quantile | 97.5% Quantile |
| Kir6.2*-GFP+SUR1<br>n = 9 | *L* | -1.05 | -1.85 | -0.45 |
|  | *D* | 0.04 | 0.00 | 0.19 |
|  | *K<sub>A</sub>* | 4.32 | 4.21 | 4.44 |
| Kir6.2*-C166S-GFP+SUR1<br>n = 6 | *L* | 0.29 | -1.04 | 1.41 |
|  | *D* | 0.84 | 0.52 | 0.95 |
|  | *K<sub>A</sub>* | 4.18 | 3.93 | 4.47 |
| Kir6.2*-GFP+SUR-K205A<br>n = 9 | *L* | -0.37 | -1.34 | 0.41 |
|  | *D* | 0.55 | 0.39 | 0.65 |
|  | *K<sub>A</sub>* | 3.76 | 3.59 | 3.95 |
| Kir6.2*-GFP+SUR-K205E<br>n = 9 | *L* | -0.18 | -1.25 | 0.70 |
|  | *D* | 0.62 | 0.42 | 0.74 |
|  | *K<sub>A</sub>* | 3.40 | 3.21 | 3.62 |
| **Single-site** |  |  |  |  |
| Kir6.2*-GFP+SUR1<br>n = 9 | *L* | -1.06 | -1.84 | -0.47 |
|  | *D* | 0.05 | 0.01 | 0.10 |
|  | *K<sub>A</sub>* | 4.33 | 4.22 | 4.44 |
| Kir6.2*-C166S-GFP+SUR1<br>n = 6 | *L* | 0.09 | -1.15 | 1.05 |
|  | *D* | 0.70 | 0.29 | 0.91 |
|  | *K<sub>A</sub>* | 4.15 | 3.88 | 4.43 |
| Kir6.2*-GFP+SUR-K205A<br>n = 9 | *L* | -0.25 | -1.30 | 0.53 |
|  | *D* | 0.18 | 0.06 | 0.32 |
|  | *K<sub>A</sub>* | 3.62 | 3.45 | 3.83 |
| Kir6.2*-GFP+SUR-K205E<br>n = 9 | *L* | -0.19 | -1.19 | 0.52 |
|  | *D* | 0.30 | 0.13 | 0.47 |
|  | *K<sub>A</sub>* | 3.31 | 3.13 | 3.50 |
| **Negative cooperativity** |  |  |  |  |
| Kir6.2*-GFP+SUR1<br>n = 9 | *L* | -0.42 | -1.38 | 0.48 |
|  | *D* | 0.15 | 0.02 | 0.29 |
|  | *K<sub>A</sub>* | 4.82 | 4.54 | 5.29 |
|  | *C* | 0.17 | 0.06 | 0.36 |
| Kir6.2*-C166S-GFP+SUR1<br>n = 6 | *L* | 0.32 | -0.96 | 1.47 |
|  | *D* | 0.83 | 0.50 | 0.94 |
|  | *K<sub>A</sub>* | 4.43 | 4.04 | 5.14 |
|  | *C* | 0.52 | 0.09 | 0.97 |
| Kir6.2*-GFP+SUR-K205A<br>n = 9 | *L* | -0.16 | -1.18 | 0.64 |
|  | *D* | 0.52 | 0.32 | 0.64 |
|  | *K<sub>A</sub>* | 4.10 | 3.73 | 4.68 |
|  | *C* | 0.35 | 0.10 | 0.91 |
| Kir6.2*-GFP+SUR-K205E<br>n = 9 | *L* | 0.03 | -1.11 | 0.99 |
|  | *D* | 0.58 | 0.32 | 0.73 |
|  | *K<sub>A</sub>* | 3.71 | 3.34 | 4.41 |
|  | *C* | 0.45 | 0.10 | 0.96 |

Our simultaneous binding and current measurements were well fit with a Monod-Wyman-Changeux (MWC)-type model (Figure 2D, E; Monod et al. (1965)) which has been previously proposed to explain K_ATP_ channel inhibition (Enkvetchakul and Nichols, 2003; Craig et al., 2008; Vedovato et al, 2015). In our MWC-type model, each ligand binding event (*K_A_*) is independent and each bound ligand favours the closed state by the same factor (*D*). Simultaneous measurement of binding (fluorescence) and gating (current) allowed us to obtain well constrained fits to our model. To obtain free parameter (*L, K_A_, D*) estimates and verify that each parameter was well and uniquely determined, we employed a Bayesian Markov chain Monte Carlo (MCMC) method previously employed by Hines et al. (Hines et al., 2014). Using this approach, we constructed posterior probability distributions for the free parameters of our MWC-type model (Figure 2F, Table 3). Based on these distributions, we estimated *K_A_* = 2.1 x 10^4^ M^−1^ (*K_D_* = 47.9 μM), *L* = 0.09 (*Ρ_open_* = 0.08), and *D* = 0.04. The very low *D* value indicates that nucleotide binding was tightly coupled to channel closure; i.e. nucleotides have a very strong preference for the closed state of the channel. The low value for *D* also explains why the channels were nearly completely inhibited at TNP-ATP concentrations at which not all the binding sites were occupied, as well as the degree to which channel inhibition is complete at saturating concentrations of TNP-ATP. Our estimate of *L* was quite low and broadly distributed. We repeated our fits with *L* fixed to a value consistent with previous single channel measurements (0.8, *P_open_* = 0.45, John et al., (1998); Enkvetchakul et al., (2000); Ribalet et al. (2006)). This had only a very small effect on our estimates of *D* and *K_A_* (Figure 2—Figure supplement 1). The broad distribution of F in our fit may represent current rundown which occurs during our patch-clamp recordings and is expected to affect the open-closed equilibrium. Cross-correlation plots (in parameter space) of the values derived from our fits produced well bounded ellipsoids, indicating that our parameters were uniquely determined (Figure 2—Figure supplement 1A).

In addition to the full MWC-type model we considered alternate models (Figure 2—Figure supplement 2). These included a model in which only the first binding event influences the open-closed equilibrium of the channel (single-binding model; Figure 2—Figure supplement 2B, Table 3), and an MWC-style model with an additional parameter ***C*** to allow for direct negative cooperativity between binding sites (negative cooperativity model; Figure 2—Figure supplement 2C, Table 3). The single-binding model yielded very similar parameter estimates to our full MWC-type model (Figure 2—Figure supplement 2D, Table 3). This is a consequence of ***D*** being so low that even in the MWC-type model most channels are closed when only a single nucleotide is bound. The cooperative model improved our fits, but not enough to justify the inclusion of an additional free parameter (see Discussion).

### Kir6.2-C166S affects the ability of bound nucleotides to close K_ATP_

To provide a rigorous test as to whether our experimental system was capable of separating nucleotide binding from subsequent channel gating, we introduced a mutation (Kir6.2-C166S) which increases *P_open_* of K_ATP_ and decreases sensitivity of the channel to block by nucleotides (Trapp et al., 1998). C166 is located near the bundle-crossing gate of Kir6.2 (Figure 3A). Other mutations at this site cause neonatal diabetes (Flanagan et al., 2006; Gloyn et al., 2006).

**Figure 3.**
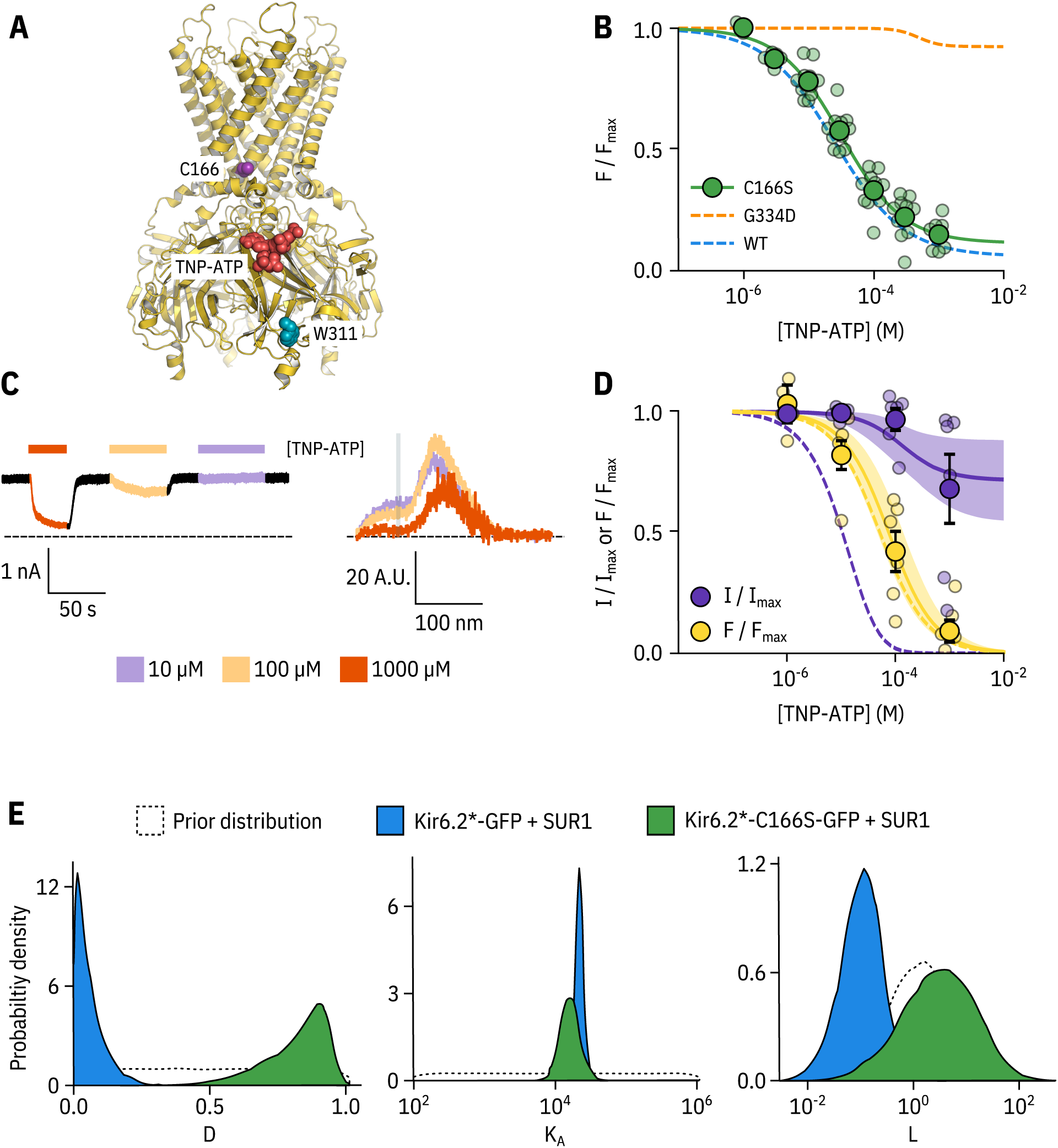
Kir6.2-C166S disrupts current inhibition, not nucleotide binding. **A.** Cartoon (from PDB accession #6BAA) showing the location of Kir6.2-C166 (purple) relative to the inhibitory nucleotide binding site (TNP-ATP from PDB accession #5XW6 shown in red), W311 is shown as blue spheres. **B.** Concentration dependence of TNP-ATP binding to unroofed membrane fragments expressing Kir6.2*-C166S-GFP + SUR1 shown in green, expressed as quenching of ANAP fluorescence. The Hill fits shown previously for Kir6.2*-GFP + SUR1 and Kir6.2*-G334D-GFP are shown in blue and orange dashed curves, respectively. Kir6.2*-C166S-GFP + SUR1: £C_50_ = 32.0 μΜ, *h =* 0.92, *E_max_ =* 0.96, n = 12, **C.** Representative current and fluorescence traces recorded simultaneously from an excised patch expressing Kir6.2*-C166S-GFP + SUR1, Exposure to different concentrations of TNP-ATP are shown by colour, **D.** Concentration-response (n = 6) for TNP-ATP inhibition of Kir6,2*-C166S-GFP + SUR1 currents (*I*/*I_max_*) and for quenching of ANAP fluorescence (*F*/*F_max_*). Data were fit with the MWC-type model. Solid curves represent the median fits and shaded areas indicate the 95% quantile intervals. Dashed curves represent the previous median fits of the MWC-type model to the Kir6.2*-GFP + SUR1 data from Figure 2D, Parameter estimates are reported in Table 3. **E.** Posterior probability distributions for the full MWC-type model fit to Kir6.2*-C166S-GFP + SUR1 or Kir6.2*-GFP + SUR1 (data from Figure 2F) overlaid on the prior probability distribution.

In unroofed membranes, Kir6.2*-C166S-GFP + SUR1 bound TNP-ATP with an *EC*_50_ very similar to that of Kir6.2*-GFP + SUR1 (Figure 3B, 32.0 μM and 25.6 μΜ, respectively), which suggests only a small change in nucleotide affinity. This is an unexpected finding, as one might expect that an increase in *P_open_* would aliosterically cause a decrease in the apparent affinity for inhibitory nucleotide binding. To resolve this conflict, we again turned to PCF (Figure 3C,D). Rundown was much slower for Kir6.2*-C166S-GFP + SUR1, which may reflect the increased *P_open_* of this construct. Measuring current inhibition in combination with nucleotide binding confirmed that whereas the apparent nucleotide affinity was unchanged by the C166S mutation, current inhibition occurred at much higher concentrations compared to binding (Figure 3D). How can we explain this paradox? Fits of the data with our MWC-type model (Figure 3D,E) suggest that, in addition to the expected effect on *L*, the C166S mutation profoundly affects the ability of bound ligand to close the channel (D) without affecting *K_A_* (Figure 3E, Table 3). We propose that, in addition to increasing the *P_open_* of the channel, C166 is also important in the transduction pathway from the inhibitory nucleotide binding site on Kir6.2 to the channel gate.

### Exploring the effect of SUR1 on nucleotide inhibition of K_ATP_

SUR1 plays a complex role in the regulation of Kir6.2. It increases the *P_open_* of the channel and allows for the activation of the channel by Mg-nucleotides (Nichols et al., 1996; Tucker et al., 1997; Babenko and Bryan, 2003; Chan et al., 2003; Fang et al, 2006). However, it also increases the sensitivity of Kir6.2 to nucleotide inhibition (Babenko and Bryan, 2003; Chan et al., 2003; Fang et al., 2006). To understand the effect of SUR1 on nucleotide inhibition of K_ATP_, we expressed Kir6.2*-GFP in the absence of SUR1 in unroofed membranes and measured TNP-ATP binding (Figure 4A). We found only a small increase (approximately 1.5-fold) in apparent *EC*_50_ compared to the same construct in the presence of SUR1 (37.6 μM and 25.6 μΜ respectively). Unfortunately, we were unable to achieve high enough expression of Kir6.2*-GFP alone to carry out PCF experiments in the absence of SUR1. However, we were able to measure currents from unlabelled Kir6.2-GFP alone (Figure 4B). As expected Kir6.2-GFP alone was much less sensitive to inhibition by TNP-ATP than Kir6.2-GFP + SUR1.

**Figure 4.**
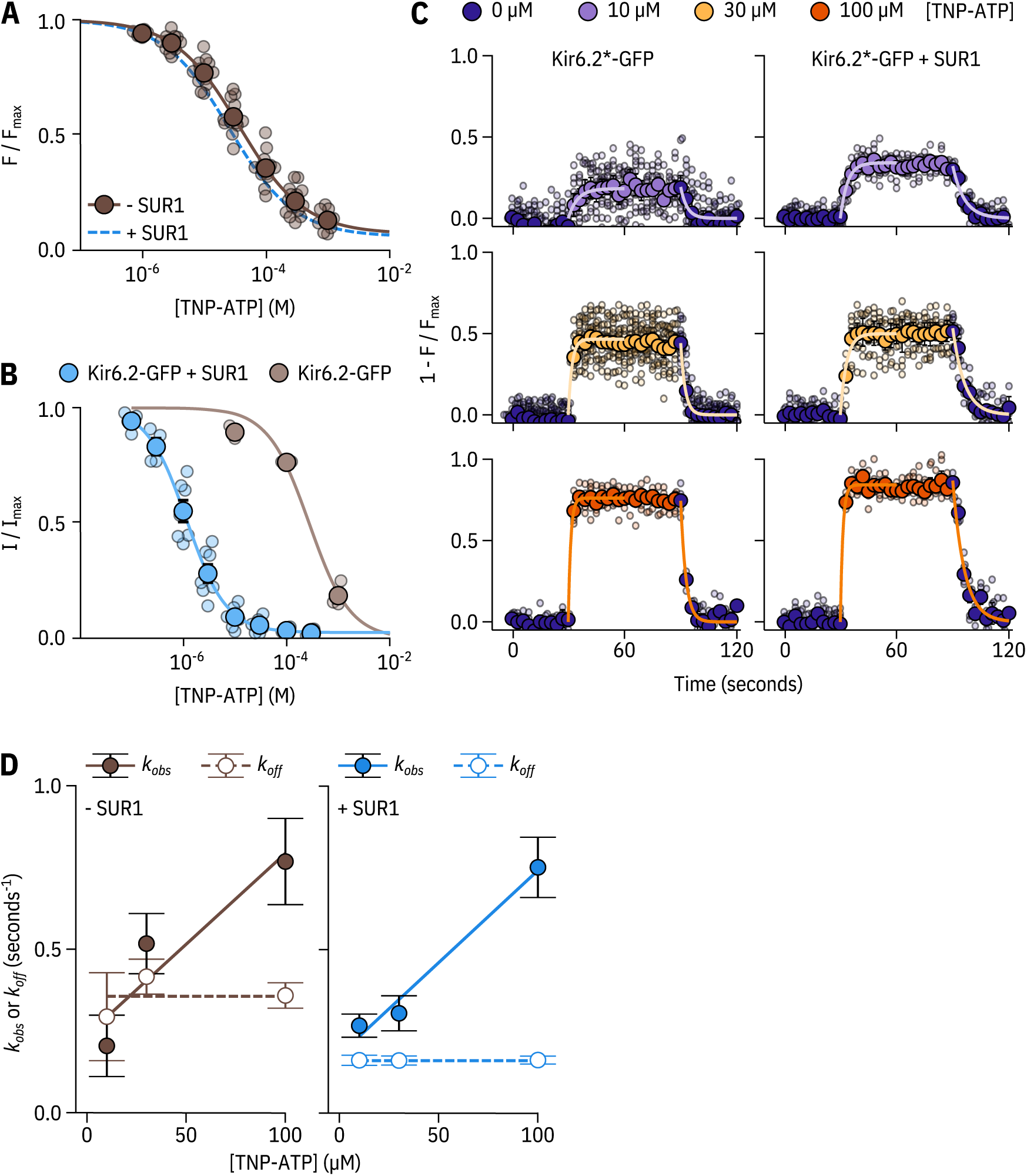
SUR.1 affects the apparent affinity for and kinetics of nucleotide binding to Kir6.2. **A.** Concentration dependence of TNP-ATP binding to unroofed membrane fragments expressing Kir6.2*-GFP without SUR1 (brown), expressed as quenching of ANAP fluorescence. The smooth curve is a descriptive Hill fit Kir6.2*-GFP (no SUR1): *EC*_50_ = 37.6 μΜ, *h =* 0.83, *E_max_ =* 0.92, n = 14. The Hill fit to Kir6.2*-GFP + SUR1 is shown as a blue dashed curve. **B.** Concentration-response curve for TNP-ATP inhibition of Kir6.2-GFP (no ANAP label) without or without co-expression of SUR1, measured in excised, inside-out patches. Kir6.2-GFP + SUR1 : *EC*_50_ *=* 1.17μΜ, *h =* 1.14, *E_max_ =* 0.97, n = 7; Kir6.2-GFP (no SUR1): *EC*_50_ *=* 273 μΜ, *h =* 1.09, *E_max_ =* 1.00, n = 3. **C.** Time-courses of TNP-ATP binding and unbinding to Kir6.2*-GFP expressed in unroofed membrane fragments in the presence or absence of SUR1. Data are displayed as 1 *-F/F_max_* so that upward deflections indicate binding and downward deflections indicate unbinding. Small data points represent individual experiments. Overlaid are larger points representing the mean ±standard error at each time point. The smooth curves are single exponential fits to the wash-on or wash-off of a given concentration of TNP-ATP. **D.** Rate constants (*k_obs_* and *k_off_*) from the exponential fits as in **C** are plotted as functions of the TNP-ATP concentration. Linear fits to were performed using equation 3. Kir6.2*-GFP (no SUR1): *k_on_ =* 5560 M^−1^ s^−1^ ±2180M^−1^ s^−1^, *k_off_ =* 0.24 s^−1^ ± 0.13 s^−1^, n = 2-6 per concentration. Kir6.2*-GFP + SUR1 : *k_on_ =* 5640 M^−1^ s^−1^ ± 812 M^−1^ s^−1^, *k_off_ =* 0.18s^−1^ ± 0.05 s^−1^, n = 3-4 per concentration. Dashed lines indicate the mean rates measured for wash-off experiments (koff) from all test concentrations combined. Kir6.2*-GFP (no SUR1): *k_off_* = 0.36s^−1^ ± 0,04s^−1^. Kir6.2*-GFP + SUR1 : *k_off_* = 0.16s^−1^ ± 0.0004s^−1^.

To determine whether SUR1 had complex effects on nucleotide binding that were not revealed in equilibrium binding experiments, we measured the time-course of TNP-ATP binding and unbinding to Kir6.2*-GFP expressed in unroofed membranes in the absence and presence of SUR 1 (Figure 4C). We fit the apparent on- (*k_obs_*) and off-rates (*k_off_*) for different concentrations of TNP-ATP to single exponential decays (equation 1). As expected, the off-rate was independent of [TNPATP]. We determined *k_on_* from the slope of linear fits to *k_obs_* as a function of nucleotide concentration, where *k_obs_* = *k_on_* * [TNPATP] + *k_off_* (Figure 4D). *k_on_* for TNP-ATP binding was nearly identical in the presence and absence of SUR1 (5641 M^−1^ s^−1^ vs. 5564M^−1^ s^−1^, respectively). However, the mean *k_off_* was roughly twice as fast in the absence of SUR1 (0.36 s^−1^ in the absence ofSURI vs. 0.16 s^−1^ in the presence of SUR1). Measuring *k_on_* and *k_off_* also provided an independent measure of *EC*_50_, ifwe assume a single-step process with *EC*_50_ given by *k_off_*/*k_on_* Using this method, we calculated the *EC*_50_ for Kir6.2*-GFP + SUR1 to be 28.3 μΜ, quite close to the 25.6 μΜ derived from steady-state measurements. The *EC*_50_ calculated from the TNP-ATP binding kinetics for Kir6.2*-GFP in the absence of SUR1 was 64.0 μΜ, higher than the value of 37.6 μΜ derived from steady-state measurements. We believe this discrepancy arises from the variability in our rate measurements. Nevertheless, the two-fold decrease in *k_off_* for TNP-ATP in the presence of SUR1 suggests that SUR1 stabilises nucleotide binding to Kir6.2. However, these binding measurements do not rule out an indirect, allosteric effect of SUR1 on nucleotide binding. To explore the effect of SUR1 more rigorously, we again turned to PCF.

As Kir6.2*-GFP expression in the absence of SUR1 was not sufficient for PCF recordings, we took a mutational approach to better understand the roieofSURI in inhibitory nucleotide binding. SUR1-K205 is located in the LO linker of SUR1, which connects the first set of transmembrane domains (TMDO) to the ABC core structure (Figure 1 A, Figure 5A; Martin et al. (2017); Puljung (2018)). This loop is adjacent to the inhibitory nucleotide-binding site on Kir6.2 and the interface between neighbouring Kir6.2 subunits. Mutations at K205 were previously shown to reduce sensitivity of K_ATP_ to nucleotide-dependent inhibition (?). Other mutations in LO are associated with neonatal diabetes (Ashcroft et al., 2017) and PHHI (Snider et al., 2013).

**Figure 5.**
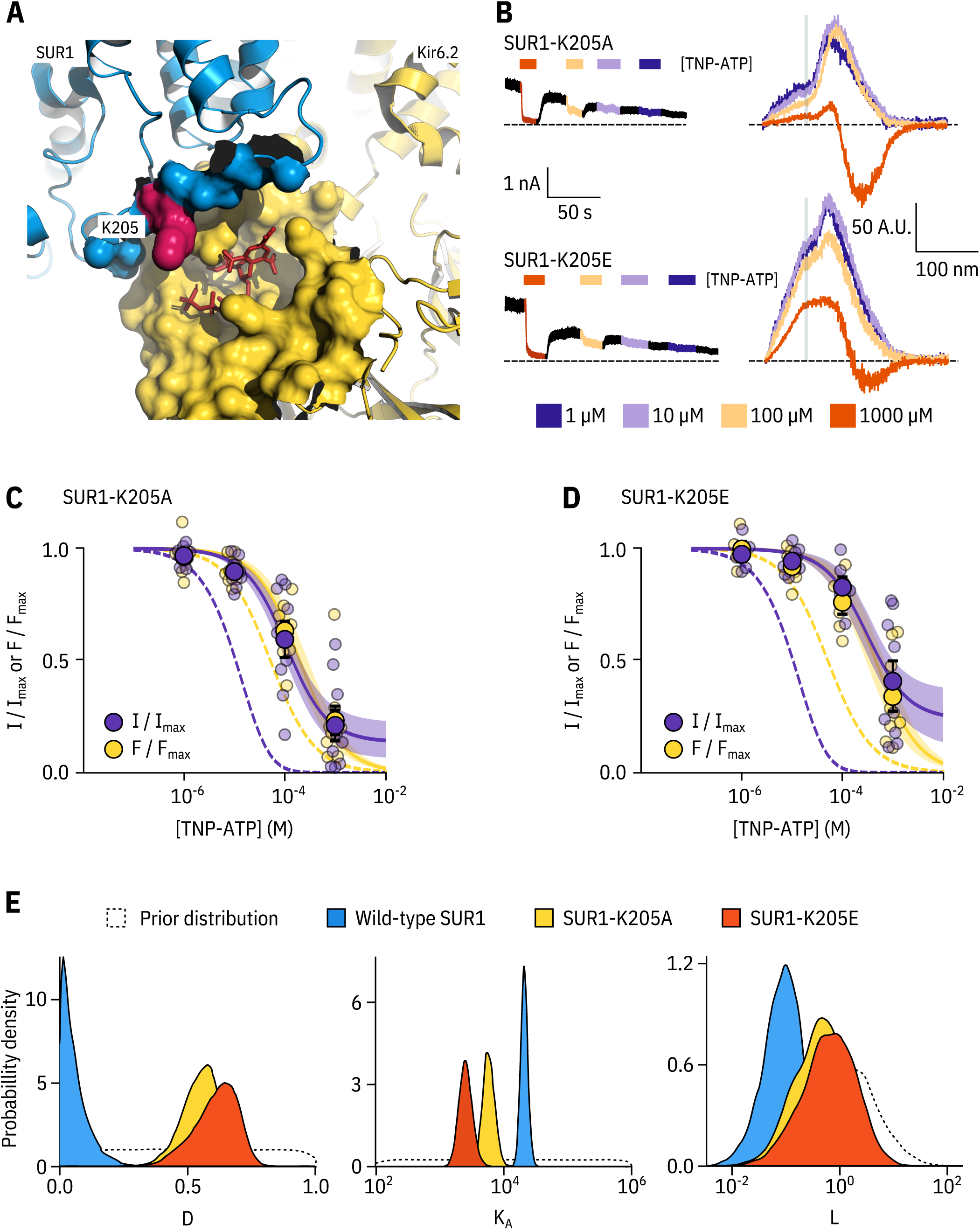
SUR1-K205 modulates both nucleotide affinity and inhibition of Kir6.2. **A.** Hydrophobic surface representation of Kir6.2 (yellow, PDB accession #6BAA) and SUR1 (blue, PDB accession #6PZI), Residue K205 on SUR1 is highlighted in pink. As this residue was built as an alanine in the structure, we used the mutagenesis tool in PyMol to insert the native lysine residue. A docked TNP-ATP molecule is shown in red, **B.** Representative current and fluorescence traces acquired simultaneously from excised patchesexpressing Kir6.2*-GFP with SUR1-K205Aor SUR1-K205E. **C,D.** Concentration-response for TNP-ATP inhibition of currents (*I*/*I_max_*) and for quenching of ANAP fluorescence (*F*/*F_max_*) in excised inside-out membrane patches expressing Kir6.2*-GFP + SUR1-K205A **(C,** n = 9) or Kir6.2*-GFP + SUR1-K205E **(D,** n = 9), Data were fit to the MWC-type model. Solid curves represent the median fits and shaded areas indicate the 95% quantile intervals. Fits to Kir6.2*-GFP + wild-type SUR1 are shown as dashed curves, **E.** Posterior probability distributions for the full MWC-type model fit to Kir6.2*-GFP co-expressed with wild-type SUR1 (fits from Figure 2), SUR1-K205A and SUR1-K205E overlaid on the prior probability distribution.

We introduced a charge neutralization (alanine, K205A) and a charge reversal (glutamate, K205E) mutation at this position and measured simultaneous nucleotide binding and current inhibition with PCF (Figure 5B,C,D). The binding and inhibition curves for TNP-ATP almost perfectly overlaid for the SUR1-K205A mutant (Figure 5C). The same was also true for SUR1-K205E (Figure 5D). Data were fit with the MWC-type model as before. Mutating K205 to an alanine or a glutamate resulted in an apparent decrease in nucleotide binding affinity (Figure 5C,D,E). This was reflected by a decrease in the estimated *K_A_* for TNP-ATP, which correlated with the degree of conservation of the mutation, i.e. we observed a larger effect for the charge reversal compared to the charge neutralization mutation (Figure 5E). However, in addition to direct effects of K205 on nucleotide binding, we also observed a shift in *D* for both mutations (Figure 5E). This suggests a dual role for SUR1 in K_ATP_ inhibition, both in contributing to nucleotide binding and in stabilizing the nucleotide-bound closed state.

## Discussion

We have developed a novel approach that allows for site-specific measurement of nucleotide binding to K_ATP_ and concomitant measurements of channel current. Performing these measurements simultaneously allowed us to examine nucleotide regulation of K_ATP_ function in great detail. We used a Bayesian approach to fit models to our combined fluorescence/current data sets to extract meaningful functional parameters with a minimum of prior assumptions. Such insights would not be possible from experiments in which macroscopic currents or binding were measured in isolation.

PCF has been used successfully by other labs to simultaneously measure ligand binding and gating in HCN channels (Biskup et al., 2007; Kusch et al., 2010; Wu et al., 2011). These groups measured fluorescence from a cyclic nucleotide analogue that increased its quantum yield when bound, minimizing background fluorescence from unbound ligand. Additional background subtraction could be performed by imaging the patches using confocal microscopy such that a region corresponding to the patch membrane could be computationally selected, thus omitting background fluorescence from the surrounding solution (Biskup et al., 2007; Kusch et al., 2010). In our PCF experiments, we used a FRET-based approach to measure ligand binding. We acquired fluorescence emission spectra, such that donor fluorescence could be separated from acceptor fluorescence by wavelength. This allowed us to directly assess binding from the quenching of donor fluorescence, which was specific to K_ATP_. FRET also provided the spatial sensitivity necessary to discriminate between nucleotide binding directly to Kir6.2 and to the nucleotide-binding sites of SUR1. We assume that any TNP-ATP bound non-specifically to our membranes would be too far from Kir6.2 to cause appreciable FRET. This assumption was confirmed by the lack of FRET between TNP-ATP and a Kir6.2*-GFP mutant (G334D), in which nucleotide binding was severely disrupted (Figure 1H).

Previous studies have suggested that K_ATP_ inhibition follows an MWC-type model (Trapp et al., 1998; Enkvetchakul and Nichols, 2003; Drain et al., 2004; Craig et al., 2008; Vedovato et al., 2015). The majority of this earlier work was performed using single-channel measurements of mutated and/or concatenated channel subunits. In this study, we confirm these results using minimally perturbed channels with nucleotide sensitivity similar to that of wild-type K_ATP_ (Figure 1—Figure supplement 2A). By using an MCMC approach to model fitting, we can also evaluate our models to assess how well the derived parameters were determined by the data. MCMC fits provide a basis for determining credible intervals for our parameter estimates. This allows for direct comparison of values derived from wild-type and different mutant constructs.

Although we did not explicitly include the effects of PIP_2_ on K_ATP_ gating in our model formulations, we assumed that the effects of PIP_2_ on *P_open_* were implicitly modelled in our parameter *L;* i.e. rundown due to dissociation of PIP_2_ manifests as a decrease in *L* rather than a change in the number of channels. Although we were able to extract identifiable parameter estimates for *L, D* and *K_A_*, our estimates of *L* for each model we considered were appreciably less well constrained than for the other parameters. We expect that this uncertainty arises from measuring a heterogeneous population of channels with regard to PIP_2_ binding. Fixing *L* to values derived from the literature (Figure 2—Figure supplement 1, Figure 3—Figure supplement 1, Figure 5—Figure supplement 1, Figure 5—Figure supplement 2) allowed us to extract estimates for *D* and *K_A_* that were functionally identical to those derived from unconstrained fits, suggesting that the uncertainty of *L* does not affect our inferences for these other parameters. Therefore, PCF represents a robust means to compare *K_A_* and *D* between different mutated K_ATP_ constructs without worrying about the confounding effects of rundown.

Previous studies suggest that, whereas K_ATP_ closure occurs via a concerted mechanism, individual nucleotide binding events at Kir6.2 are not equivalent (Markworth et al., 2000). Earlier attempts to determine the stoichiometry of inhibitory nucleotide binding to Kir6.2(i.e. how many ATPs must bind to induce channel closure) have produced models ranging from those in which binding of a single nucleotide completely shuts K_ATP_ to an MWC-type model in which each binding event is independent and contributes equally to channel closure (Trapp et al., 1998; Markworth et al., 2000; Enkvetchakul and Nichols, 2003; Drain et al., 2004; Wang et al., 2007; Craig et al., 2008; Vedovato et al., 2015). To resolve this controversy, we fit our data with both single-binding and MWC-type models. At very low values for *D*, such as we derived from our experiments, the predictions of both models are functionally very similar. Even in our MWC-type model, we expect most K_ATP_ channels to be closed when just one molecule of nucleotide is bound.

It has been proposed that there is direct negative cooperativity between binding events at different subunits on Kir6.2 (Wong et al., 2007). We fit our data to an extended MWC-type model including an additional free parameter (C), representing negative binding cooperativity between subunits (Figure 2—Figure supplement 2). Not surprisingly this model improved the fit to our data as assessed by the Bayes factor, which represents the marginal likelihood of one model over another to explain our observations (Wagenmakers, 2007; Gronau et al., 2017)*. We* also tested the cooperative model using approximate leave-one-out cross validation, which assesses the ability of a model to predict new or out-of-sample data using in-sample fits. Although in this work, we are primarily concerned with the inferences made from our fits, the ability of a model to make predictions is a good measure of its usefulness. Based on this criterion, the cooperative model has no more predictive accuracy than either the MWC-type model or the single-binding model. Therefore, the inclusion of an additional free parameter is not justified. Furthermore, whereas the cooperative model yielded good fits with identifiable parameters for Kir6.2*-GFP + SUR1 channels, it failed to do so for all the mutants considered. Thus, this model did not allow for direct comparison between constructs. However, it remains a possibility that these mutations function in part by abolishing binding cooperativity between subunits.

We performed all our experiments on mutated, tagged channels using a fluorescent derivative of ATP. This allowed us to fit mechanistic models and readily compare between mutated constructs that affect nucleotide inhibition of K_ATP_. This raises an obvious question: how relevant are our findings to inhibition of wild-type K_ATP_ by ATP? In a previous paper, we estimated *D* and *K_A_* from an MWC-type model based on fits to published data for ATP inhibition of wild-type Kir6.2 + SUR1 (Proks et al., 2010; Vedovato et al., 2015). The value we obtained for *D* (0.03) was quite similar to that we report here from our PCF measurements (0.04). We also obtained a similar estimate for *K_A_* in our previous model (3.0 x 10^4^M^−1^ vs 2.1 x 10^4^M^−1^ from our PCF experiments). Despite obtaining similar parameters, past experiments in which only ionic currents were measured, did not allow us to distinguish between competing gating models. Measuring currents and fluorescence simultaneously allowed for better model selection and aided in our ability to identify constrained parameters.

We compared the parameters derived for inhibitory nucleotide binding to those estimated for nucleotide activation of K_ATP_ based on experiments in which currents and binding were measured in separate preparations (Puljung et al., 2019). In those experiments, we estimated a value for *E*, the factor by which binding of MgTNP-ADPtoSUR1 stabilized channel opening, of 2.2. Although this value was derived using a different nucleotide, it still provides an approximate basis for comparing the coupling of nucleotide stimulation through SUR1 to nucleotide inhibition via binding to Kir6.2. If both activation and inhibition proceed via MWC-type models, the open closed equilibrium at saturating nucleotide concentrations is given by *L* multiplied by *E*^4^ or *E*^4^, respectively. The degree of stabilization of the open state of K_ATP_ can be calculated as −*RT* In *E*^4^ for activation. Stabilization of the closed state is given by −*RT* In *D*^4^. Based on our observations, saturating concentrations of MgTNP-ADP stabilized the open state by −1.9kcal mol^−1^ (−7.9 kJ mol^−1^). At saturating concentrations, TNP-ATP stabilized the closed state of K_ATP_ by −7.6 kcal mol^−1^ (31.8 kJ mol^−1^). Thus, at conditions under which both excitatory and inhibitory nucleotide binding sites are saturated, inhibition dominates, which is consistent with published measurements of wild-type K_ATP_ in the presence of Mg^2+^ (). In our previous study, we estimated *K_A_* for MgTNP-ADP binding to the stimulatory second nucleotide binding site of SURI to be 5.8×10^4^M^−1^ (*K_D_* = 17μΜ), higher affinity than the *K_A_* we report here for TNP-ATP binding to the inhibitory site on Kir6.2 (2.1 × 10^4^M^−1^,Æ_p_ =48μΜ). Higher affinity binding to the stimulatory site may explain the ability of MgADP to increase K_ATP_ currents in the presence of ATP (Gribble et al., 1998). This phenomenon may also explain the bell-shaped MgADP concentration-response curve for K_ATP_, which shows an increase in current at low concentrations, followed by inhibition at higher concentrations (Proks et al., 2010; Vedovato et al., 2015). Future experiments in which activation and inhibition are measured by PCF for the same ligand will allow us to model the complex response of K_ATP_ under conditions where all three nucleotide binding sites simultaneously affect channel gating (i.e. in the presence of Mg^2+^).

Mutations that cause neonatal diabetes reduce the sensitivity of K_ATP_ to nucleotide inhibition, and reduction in nucleotide sensitivity is broadly correlated with disease severity (McTaggart et al., 2010)*. We* studied two residues on Kir6.2 that have been implicated in diabetes and have been proposed to affect nucleotide sensitivity via different mechanisms. We find that G334D drastically reduced the apparent affinity for nucleotide binding to K_ATP_ in unroofed membranes. In our MWC-type models, this could only be explained by a dramatic decrease in *K_A_.* This corroborates earlier hypotheses that mutating G334 directly disrupts inhibitory nucleotide binding to Kir6.2 (Drain et al., 1998). Due to poor expression, we were unable to test this construct using PCF. Therefore, we could not obtain accurate estimates of *K_A_* and *D*.

In contrast to G334D, the C166S mutation does not directly affect nucleotide binding to Kir6.2, but rather disrupts the ability of bound nucleotide to close the channel. This contributes to the decreased nucleotide sensitivity which was previously attributed solely to an increased *P_open_.* In the future, we hope to use this rigorous approach to assess a whole panel of neonatal diabetes mutations in Kir6.2to better understand the mechanism by which they cause disease.

Using PCF allowed us to probe more deeply into the role of SUR1 in regulating nucleotide inhibition of K_ATP_. The cytoplasmic LO loop of SURI was previously implicated in modulation of *P_open_* and nucleotide sensitivity of Kir6.2 (Babenko and Bryan, 2003; Chan et al., 2003; Pratt et al., 2012). We find that, in addition to directly contributing to tighter nucleotide binding at Kir6.2, SUR1 plays a critical role in preferentially stabilising the closed state of the channel when nucleotides are bound. Whereas a single nucleotide-binding event is sufficient for channel closure when Kir6.2 is associated with wild-type SUR 1, mutating residue K205 reduced the ability of a single nucleotide to close the channel. This difference manifests in both our MWC-type and single-binding models.

In addition to providing mechanistic insights into disease-associated mutations in Kir6.2, our PCF-based approach allows us to probe the interactions between Kir6.2 and SUR1 on two different levels. As we show here, we can use this method to examine the effects of SUR1 on inhibitory nucleotide binding to Kir6.2. We can also adapt this method to study activation of Kir6.2 by nucleotides bound to the stimulatory sites on SUR1. Mutations in SUR1 that cause neonatal diabetes may do so by disrupting inhibitory binding/gating or enhancing the stimulatory effects of nucleotides. The formalism developed in this study provides a rigorous way to mechanistically assess the effects of these mutations. Our approach should be readily adaptable to the study of other nucleotide-gated channels including the cystic fibrosis transmembrane conductance regulator (CFTR, also an ABC-family protein) and purinergic P2X receptors.

## Materials and Methods

### Key resources table

Key resources table.
| Reagent type (species) or resource | Designation | Source or reference | Identifiers |
| --- | --- | --- | --- |
| Cell line | HEK-293T | LGC Standards (ATCC CRL-3216) |  |
| Transfected construct (*Escherichia coli*) | pANAP | Addgene |  |
| Transfected construct | pcDNA4/TO | Addgene |  |
| Transfected construct (*Aequorea victoria*) | pCGFP_EU | Gouaux Laboratory (Vollum Institute, Portland OR USA) |  |
| Transfected construct (*Homo sapiens*) | peRF1-E55D | Chin Laboratory (MRC Laboratory of Molecular Biology, Cambridge UK) |  |
| Antibody | Anti-HA High Affinity; Rat monoclonal antibody (clone 3F10) | Roche | (Roche Cat# 11867423001, RRID:AB_10094468) |
| Antibody | Peroxidase-AffiniPure Goat Anti-Rat IgG (H + L) antibody | Jackson ImmunoResearch Labs | (Jackson ImmunoResearch Labs Cat# 112-035-003, RRID:AB_2338128) |
| Chemical compound, drug | trinitrophenyl-ATP (TNP-ATP) | Jena Bioscience (Jena, Germany) |  |
| Chemical compound, drug | L-3-(6-acetylnaphthalen-2-ylamino)-2-aminopropionic acid | Asis Chemicals (Waltham, MA) |  |

### Molecular biology

Human Kir6.2 and SUR1 were subcloned into pcDNA4/TO and pCGFP_EU vectors for expression of wild-type and GFP-tagged constructs, respectively. pcDNA4/TO and pANAP were obtained from Addgene. peRF1-E55D and pCGFP_EU were kind gifts from the Chin Laboratory (MRC Laboratory of Molecular Biology, Cambridge, UK) and the Gouaux Laboratory (Vollum Institute, Oregon, USA) respectively. Amber stop codons and point mutations were introduced using the QuikChange XL system (Stratagene; San Diego, CA). All constructs were confirmed by DNA sequencing (DNA Sequencing and Services, University of Dundee, Scotland).

### Cell culture and channel expression

HEK-293T cells were obtained from and verified/tested for mycoplasma by LGC standards (ATTC CRL-3216, Middlesex, UK). Our working stock tested negative for mycoplasma contamination using the MycoAlert Mycoplasma Detection Kit (Lonza Bioscience; Burton on Trent, UK). Cells were plated onto either poly-L-lysine coated borosilicate glass coverslips (VWR International; Radnor, PA) or poly-D-iysine coated glass-bottomed FluoroDishes(FD35-PDL-100, World Precision Instruments). ANAP-tagged Kir6.2 constructs were labelled using amber stop codon suppression as described by Chatterjee et al (Chatterjee et al., 2013). Transfections were carried out 24 hours after plating using TranslT-LT1 (Mirus Bio LLC; Madison, Wl) at a ratio of 3 pi per pg of DNA. Unless specified otherwise, all transfections included a Kir6.2 construct with an amber stop codon (TAG) at position 311 (Kir6.2-W311^TAG^), SUR1, pANAP and eRF1-E55D in the ratio 0.5:1.5:1:1. Transfected cells cultured in Dulbecco’s Modified Eagle Medium (Sigma; St. Louis, MO) + 10% foetal bovine serum, 100 U ml^−1^ penicillin and lOOpgml’^1^ streptomycin (Thermo Fisher Scientific; Waltham, MA) supplemented with 20 mM ANAP (free acid, AsisChem; Waltham, MA). Ceils were incubated at 33 °C and in the presence of 300 μΜ tolbutamide to enhance protein expression and channel trafficking to the plasma membrane (Yan et al., 2007; Lin et al., 2015). eRF1-E55D was included to increase efficiency of ANAP incorporation (Schmied et al., 2014). Experiments were carried out 2-4 days after transfection. We also expressed constructs labelled with ANAP at positions 1182, F183, F198, and 1210. Kir6.2-F183*, Kir6.2-F198*, and Kir6.2-l210* co-expressed with SUR1 did not produce sufficient currents for subsequent experimentation. Mutations at 1182 are known to produce profound effects on nucleotide inhibition of K_ATP_ (Li et al., 2000). Thus, we did not consider this site for further experimentation.

### Western blots

Transfected HEK-293T ceils grown in 6-well plates were harvested in cold PBS (Life Technologies Limited; Paisley, UK), pelleted at 0.2 x g for 2.5 minutes and resuspended in lysis buffer containing 0.5% Triton X-100, 100mM potassium acetate, and a complete protease inhibitor tablet (1 tablet/SQml, Roche; Basel, Switzerland), buffered to pH 7.4. After a 30-minute benzonase (Sigma) treatment at room temperature, samples were mixed with a DTT containing reducing agent and loading buffer (NuPAGE, Invitrogen; Carlsbad, CA)and run on a precast Bis-Tris 4-12% poly-acrylamide gel at 200V for 40 minutes. Proteins were wet transferred overnight onto polyvinylidene difluoride (PVDF) membranes (Immobilon P, Merck Millipore; Burlington, VT) in 25mM Tris, 192mM glycine, 20% methanol, and 0.1% SDS at 10V on ice. Membranes were blocked with 5% milk in TBS-Tw (150mM NaCI, 0.05% Tween 20, 25 mM mM Tris, pH 7.2) before staining for 30 minutes with a 1:1000 dilution of rat anti-HA monoclonal antibody in TBS-Tw (clone 3F10, Roche). After washing with TBS-Tw, membranes were incubated for 30 minutes with a 1:20,000 dilution of HRP-conjugated goat anti-rat polyclonal antibodies in TBS-Tw (jackson ImmunoResearch; Ely, UK). Detection was performed using the SuperSignal West Pico Chemiluminescent Substrate (Thermo Fisher) and a C-DiGit Blot Scanner (Licor Biosciences; Lincoln, NE). Analysis was performed using custom code written in Python.

To confirm our ability to express full-length Kir6.2*-GFP, we performed western blots for HA-tagged Kir6.2 constructs in detergent-solubilized HEK-293T cells (Figure 1—Figure supplement 1C). The HA tag plus a short linker (YAYMEKGITDLAYPYDVPDY) was inserted in the extracellular region following helix M1 of Kir6.2 between L100 and A101. Transfection of wild-type Kir6.2-HAor Kir6.2-HA-GFP resulted in two bands on the western blots. The upper bands were close to the expected sizes for full-length Kir6.2-HA and Kir6.2-HA-GFP (46 kDa and 77 kDa, respectively).

We consistently observed a lower molecular weight band as well. This band must correspond to an N-terminally truncated Kir6.2 product, as the apparent molecular weight shifted with addition of the C-terminal GFP tag. Based on the molecular weight, we predict that the truncated protein product initiated from a start codon in the first transmembrane domain. Therefore, we believe it is unlikely that this protein would form functional channels or traffic to the plasma membrane. When Kir6.2-W31 1^tag^-HA or Kir6.2-W31 1^tag^-HA-GFP were co-transfected with SUR1, pANAP, and eRF1-E55D, and cel Is were cultured in the presence of ANAP, the western blots were similar to wild-type Kir6.2-HA or Kir6.2-HA-GFP. Over 90% full-length Kir6.2*-HA-GFP was produced under these conditions (Figure 1—Figure supplement 1D). We were unable to quantify the percentage of full-length Kir6.2*-HA produced as the C-terminally truncated band resulting from termination at the TAG codon was very similar in size to the N-terminally truncated band. Co-expression with SUR1 increased the percentage of full-length Kir6.2*-HA-GFP produced (Figure 1—Figure supplement 1D). In the absence of ANAP, we did not observe any full-length Kir6.2, indicating that there was no read-through of the amber (TAG) stop codon (Figure 1—Figure supplement 1D).

### Confocal microscopy

Confocal imaging was performed using a spinning-disk system (Ultra-VIEWVoX, Perkin Elmer; Waltham, MA) mounted on an 1X31 microscope (Olympus; Southend-on-Sea, UK) with a Plan Apo 60x oil immersion objective (NA = 1.4), provided by the Micron Advanced Bioimaging Unit, Oxford. Transfected HEK-293T cells were incubated for 15 minutes with 1 nM CellMask Deep Red (Thermo Fisher) to stain plasma membranes before washing with PBS and imaging. ANAP was excited with a solid-state laser at 405 nM. GFP and CellMask were excited with an argon laser at 488 nM and 633 nM respectively. Images were captured on an EMCCD camera (ImagEM; Hamamatsu Photonics; Welwyn Garden City, UK) binned at 2 x 2 pixels and analysed using Python. A median filter with a box size of 32 x 32 pixels was applied to improve the signal-to-noise ratio by reducing background fluorescence.

We examined the surface expression of our ANAP-labelled constructs using confocal microscopy (Figure 1 —Figure supplement 1 A,B). When Kir6.2-W311 ^tag^-GFP was co-transfected with SUR1 along with pANAP and eRF1-E55D in the presence of ANAP, the ANAP and GFP fluorescence were colocalized at the plasma membrane. When wild-type Kir6.2-GFP was transfected under the same conditions, only GFP fluorescence was observed at the plasma membrane. ANAP fluorescence was diffuse and confined to the cytoplasm or intracellular structures. Thus, the plasma-membrane ANAP signal was specific for Kir6.2*-GFP.

### Surface expression assays

We measured surface expression of HA-tagged Kir6.2 subunits using an approach outlined by Zerangue et al. (Zerangue et al., 1999; Puljung et al., 2019). Ceils were plated on 19 mm coverslips coated with poly-l-lysine and transfected as described above. Following incubation, cells were rinsed with PBS before fixation with 10% formalin for 30 minutes at room temperature. After washing again, ceils were blocked with 1% BSA in PBS for 30 minutes at 4°C before a 1-hour incubation at 4°C with a 1:1000 dilution (in PBS) of rat anti-HA monoclonal antibodies. Cells were then washed 5 times on ice with 1% BSA in PBS followed by a 30-minute incubation at 4 °C with a 1:2000 dilution of HRP-conjugated goat anti-rat polyclonal antibodies. Cells were washed 5 times in PBS+ 1% BSA and 4times in PBS. Coverslips were removed from the culture dishes and placed in clean, untreated dishes for measurement. 300 μ I of Super Signal ELISA Femto Maximum Sensitivity Substrate (Thermo Fisher) was added to each sample and the luminescence was measured using a Glomax 20/20 Luminometer (Promega; Madison, Wl) after a 10 second incubation.

HEK-293T cells were transfected with Kir6.2 constructs with or without a TAG stop codon corresponding to position 311. Cells were co-transfected with pANAP and eRF1-E55D in the presence or absence of SUR1 and cultured with or without ANAP. Wild-type Kir6.2-HA and Kir6.2-HA-GFP in the presence of SUR1 were included as positive controls. Kir6.2 constructs with no HA tag served as negative controls. In the presence of ANAP, we observed strong trafficking of Kir6.2*-HA-GFP to the plasma membrane, but much less trafficking of Kir6.2*-HA (Figure 1—Figure supplement 1E). When cells were cultured in the absence of ANAP, we observed little to no Kir6.2 surface expression from cells that were transfected with Kir6.2-W311^TAG^-HA or Kir6.2-W31 1^tag^-HA-GFP suggesting that prematurely truncated constructs did not traffic to the plasma membrane. In the absence of SUR1, surface expression was weak for both wild-type and tagged constructs, despite the reported ability of Kir6.2-GFP to traffic to the plasma membrane in the absence of SUR1 (John et al., 1998; Makhina and Nichols, 1998).

### Epifluorescence imaging and spectroscopy

Epifluorescence imaging and spectroscopy were performed using a Nikon Eclipse TE2000-U microscope with a 60x water immersion objective (Plan Apo VC, NA =1.2, Nikon; Kingston upon Thames, UK) or a 100xoil immersion objective (Nikon, Apo TIRF, NA = 1.49). Imaging of ANAP was performed using a 385 nm LED source (ThorLabs; Newton, NJ) with a 390/18 nm band-pass excitation filter, an MD416dichroic and a 479/40 nm band-pass emission filter (all from ThorLabs). GFP was imaged using a 490 nm LED source (ThorLabs) with a 480/40 nm band-pass excitation filter, a DM505 dichroic, and a 510nm long-pass emission filter (all from Chroma; Bellows Falls, VT). Fluorescence spectra were collected by exciting ANAP as above but using a 400 nm long-pass emission filter (ThorLabs), then passing emitted light through an IsoPlane 160 Spectrometer (Princeton Instruments; Trenton, NJ) with a 300 g mm^−1^ grating. Images were collected with 0.1 s to 1 s exposures on a Pixis 400BR_eXcelon CCD (Princeton Instruments).

### Electrophysiology

Patch pipettes were pulled from thick-walled borosilicate glass capillaries (GC150F-15, Harvard Apparatus; Holliston, MA) to a resistance of 1.5 ΜΩ to 2.5 ΜΩ when filled with pipette solution. Currents were recorded at −60 mV from excised inside-out patches using an Axopatch 200B amplifier equipped with a Digidata 1322A digitizer and using pClamp 10 software (Molecular Devices; San Jose, CA). Currents were low-pass filtered at 5 kHz and digitized at 20 kHz. The bath solution (intracellular) contained 140 mM KCI, 10 mM HEPES, 1 mM EDTA and 1 mM EGTA (pH 7.3 with KOH). The pipette solution (extracellular) contained 140mM KCI, 10 mM HEPES and 1 mM mM EDTA (pH 7.4 with KOH). All experiments were carried out in Mg^2+^-free conditions. Currents were leak corrected using the current remaining in bath solution containing 5 mM barium acetate at 60 mV, assuming a linear leak with a reversal potential of OmV. Inhibition was calculated and corrected for rundown by alternating test concentrations of nucleotide solution with nucieotide-free solution, then expressing the test currents as a fraction of the average of the control currents before and after the test solution as described previously (Proks et al, 2010).

### Unroofed binding measurements

Unroofed membranes were prepared as described previously (Heuser, 2000; Zagotta et al., 2016; Puljung et al., 2019). A coverslip plated with transfected HEK-293T cells was removed from the culture media and rinsed with PBS. The coverslip was then briefly sonicated using a probe sonicator (Vibra-cell; Newtown, CT) leaving behind adherent plasma membrane fragments. Cells cultured on FluoroDishes were rinsed and sonicated directly in the dish. Unroofed membrane fragments were nearly invisible in bright-field images and identified by their GFP and ANAP fluorescence. Fluorescent TNP-nucleotides (Jena Bioscience; Jena, Germany) were diluted in bath solution and perfused onto unroofed membranes using a valve controlled microvolume superfusion system (pFlow, ALA Scientific Instruments; Farmingdale, NY).

Fluorescence spectra were collected as described above. A region of interest corresponding to the membrane fragment was manually selected and line-averaged for each wavelength. A similarly sized region of background was selected and averaged, then subtracted from the spectrum of interest. After subtraction, ANAP intensity was calculated by averaging the fluorescence intensity measured between 469.5 nm and 474.5 nm. Bleaching was corrected by fitting the normalised ANAP intensity of exposures taken during perfusion with nucleotide-free solution to a single exponential decay of the form

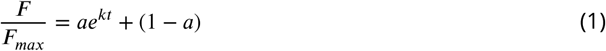

then using the fit to correct the intensity of exposures taken during perfusion with test nucleotide solutions.

For kinetic measurements, the solution changer and camera were controlled using pClamp 10 software coupled to a Digidata 1322A digitizer. Each fragment of unroofed membrane was exposed three times to the same test concentration of nucleotide. Spectra were acquired every three seconds. These technical replicates were averaged and presented as a single experiment. Bleaching was corrected by fitting the ANAP intensity of the last ten spectra acquired during each nucleotide-free solution wash to equation 1.

### Patch-ciamp fluorometry

The tip of the patch pipette was centred on the slit of the spectrometer immediately after patch excision. Currents were measured as described above. Fluorescence emission spectra from the excised patch were acquired concurrently with current measurements, both during test solution application as well as nucleotide-free solution. Background subtraction was slightly imperfect due to the exclusion of TNP-ATP from volume of the glass of the pipette, resulting in spectra that have negative intensities at the TNP-ATP peak at high nucleotide concentrations. However, this over-subtraction does not affect the size of the ANAP peak, which we used to quantify nucleotide binding.

### Data processing and presentation

Raw spectrographic images and current traces were pre-processed in Python and Clampfit (Axon) before analysis with R. Where applicable, all experimental data points are displayed in each figure. The number of experiments is reported in the figure legends and tables. To help visualise uncertainty and prevent some data points being hidden, they are arranged with a small amount of horizontal jitter; vertical position remains unaffected. Unless otherwise stated, summary statistics are overlaid as the mean with error bars representing the standard error of the mean. Where these error bars are not visible, they are smaller than the size of the point used for the mean.

Hill fits to fluorescence quenching were nonlinear least-squares fits to the following equation:

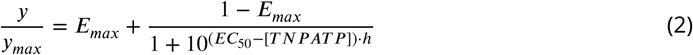

where *y* represents normalised fluorescence intensity and *EC*_50_ and [*TNPATP*] are log_10_ values. Current inhibition data were fit to the same equation but with *y* representing normalised current magnitude, *IC*_50_ instead of *EC*_50_, and *I_max_* instead of *E_max_*.

*k_obs_* values from single exponential fits with equation 1 to the wash-on and wash-off of TNP-ATP in the time-course experiments were fit with the linear equation:

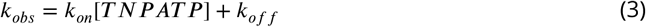

The gradient of the linear fit to the observed on-rate (*k_obs_*) is equivalent to *k_on_ ; k_off_* is the intercept at zero [*TNPATP*]. We also measured *k_off_* directly from the dequenching of ANAP following TNPATP wash-off. As expected, these values were independent of the [*TNPATP*] applied.

### Bayesian model fitting

The MWC-type models considered (Figure 2 and Figure 2—Figure supplement 2) were formulated as follows:

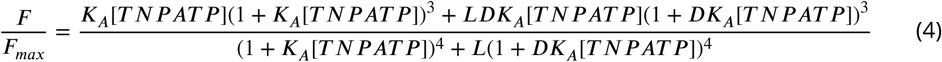

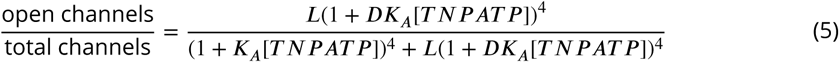

When no ligand is present (i.e. when [*TNPATP*] = 0), equation 5 becomes:

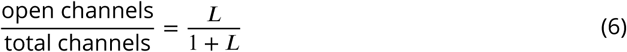

We can use this to normalise the predicted changes in the open fraction to an observed change in current as:

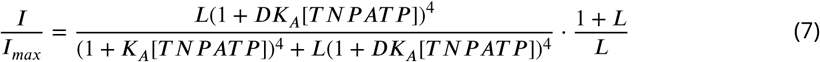

Two variations on the full MWC model were also considered, and diagrammatic formulations are shown in Figure 2 - Figure supplement 1. The first was similar to the MWC-type model, except that the channels close after one molecule of TNP-ATP binding with subsequent binding events having no effect.

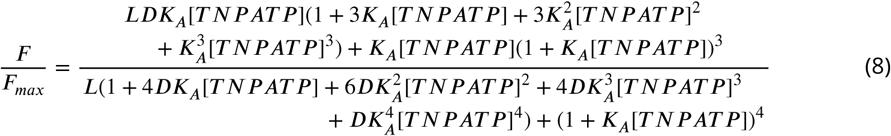

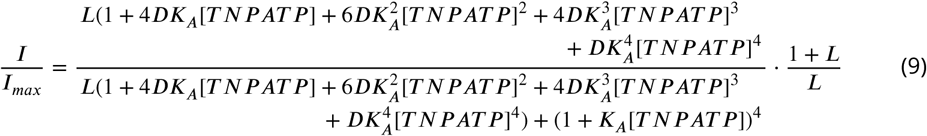

The second alternate model was the same as the full MWC model, but with an additional term ***C*** describing binding co-operativity between Kir6.2 subunits.

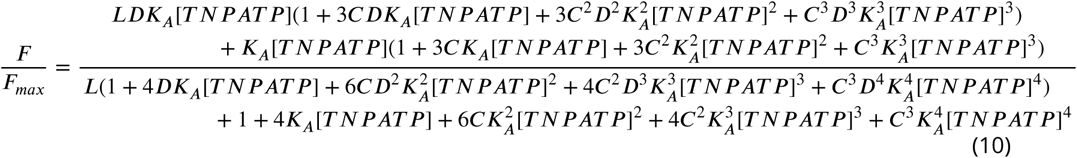

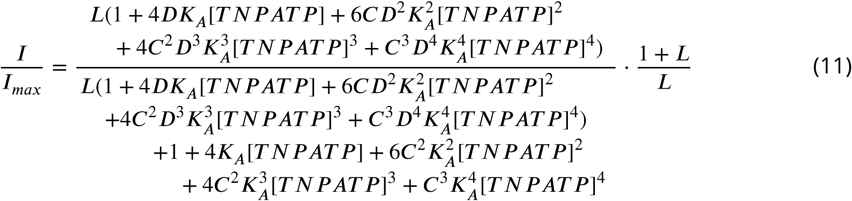

Each model was fit to the combined patch-clamp fluorometry datasets using the brms package (Gelman et al., 2015; Burkner, 2017) in R. Prior probability distributions for each parameter were supplied as:

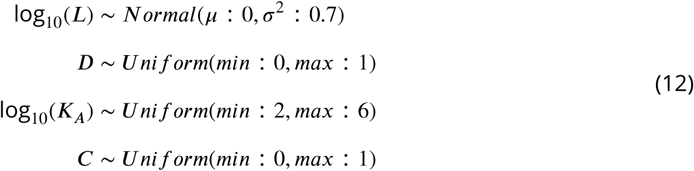

so that all priors are flat apart from L, which is weakly informative with 99% of its density falling between unliganded open probabilities of 0.01 and 0.99, and 85% falling between 0.1 and 0.9.

Each model was run with 4 independent chains for 10,000 iterations each after a burn-in period of 20,000 iterations, saving every 10th sample for a total of 4,000 samples per model. Each model parameter achieved a minimum effective sample size of 3,500 and a potential scale reduction statistic (R̂) of 1.00. Where applicable, the posterior probabilities of each parameter are reported as the median and the 95% equal-tai led interval. Bayes factors were calculated using bridge-sampling (Gronau et al., 2017), and leave-one-out cross-validation (LOO-CV) was performed using the loo package (Vehtari et al., 2017).

### Docking

Computational docking of TNP-ATP into the nucleotide binding site of Kir6.2 was performed using AutoDock-Vina (Trott and Olson, 2010) and Pymol (Schrödinger, LLC; New York, NY). 11 TNP-ATP structures from the Protein Data Bank (PDB accession #s 1I5D, 3AR7, 5NCQ, 5SVQ, 5XW6, 2GVD, 5A3S, 2PMK, and 3B5J) were used as starting poses and a 15×11.25×15 Å box was centred on the ATP bound to Kir6.2 in PDB accession #6BAA (Martin et al., 2017). Protonation states for each residue were assigned using PDB2PQR and PROPKA 3.0 (Dolinsky et al., 2004). The modal highest-scoring pose from the docking run was selected (PDB accession #5XW6, Kasuya et al. (2017)) and distances were measured from a pseudo atom at the centre of the fluorescent moiety. TNP-ATP (PDB #3AR7, Toyoshima et al. (2011)) was positioned into the first nucleotide binding domain of SUR1 (PDB #6PZI, Martin et al. (2019)) using the alignment tool in Pymol.

### Chemicals and stock solutions

Unless otherwise noted, ail chemicals were obtained from Sigma. TNP-ATP was obtained as a 10 mM aqueous stock from Jena Bioscience and stored at −20 °C. 1 mM aqueous stocks of ANAP-TFA were prepared by dissolving the free acid in 30 mM NaOH, and were stored at −20°C. Tolbutamide stocks (50 mM) were prepared in 100 mM KOH and stored at −20 °C.

### Data availability

All data sets and the code used to analyse and present them are available on GitHub at https://github.com/smusher/KATP_paper_2019

## Acknowledgments

We wish to thank Raul Terron Exposito for technical assistance and Dr. Natascia Vedovato for helpful discussions. James Cantley provided access to the Licor scanner for western blots. This work was supported by the Biotechnology and Biological Science Research Council (BB/R002517/1) and the Wellcome Trust Oxion graduate program.

---

**Figure 1 - figure supplement 1.**
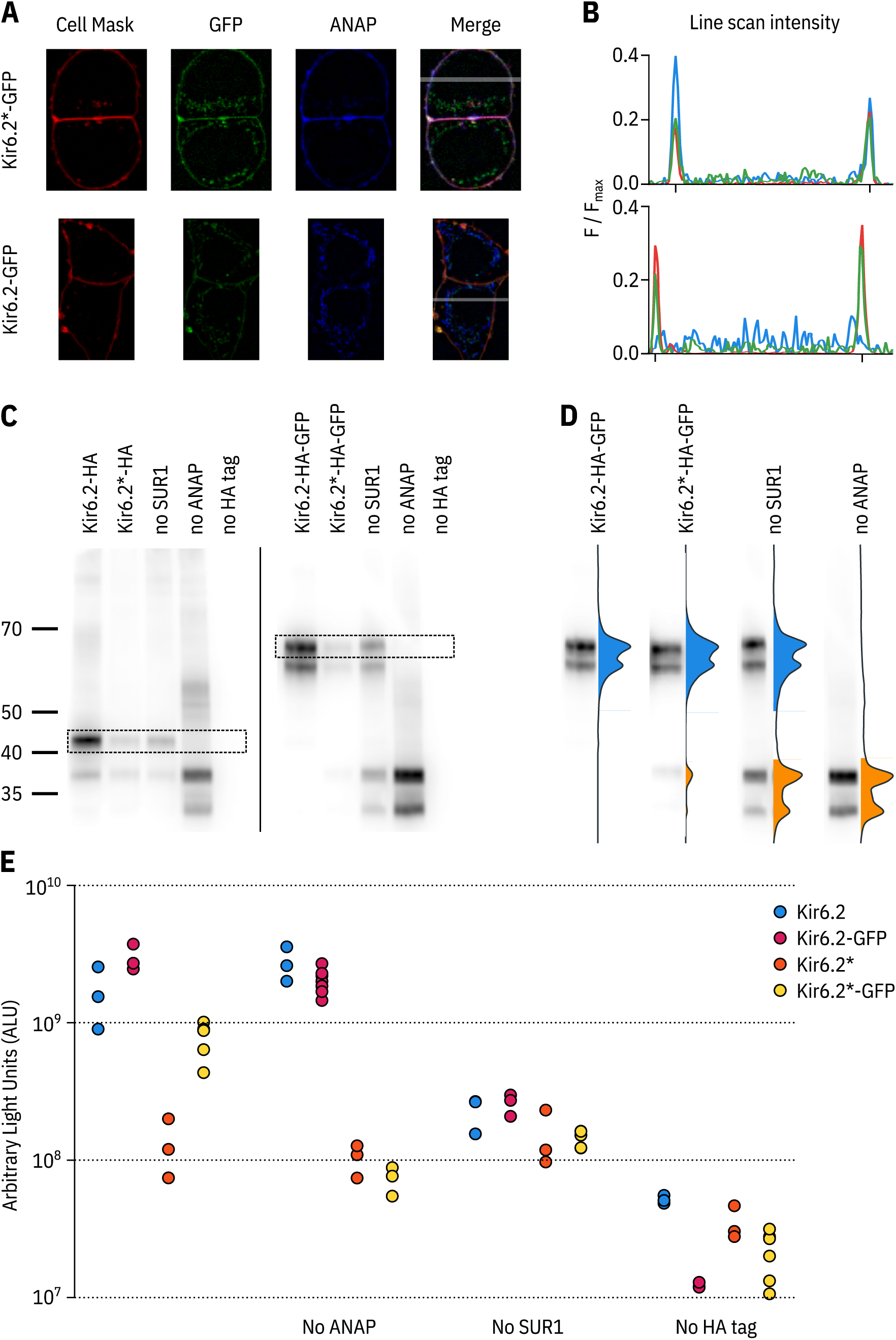
ANAP labelling is specific and only full-length Kir6.2 is expressed at the cell membrane. **A.** Confocal images of HEK-293T cel Is transfected with Kir6.2*-GFP + SUR1 (top panel) or Kir6.2-GFP + SUR1 (bottom panel). Cells were stained with Cell Mask Deep Red to label the plasma membrane. The grey band in the merged image is a 5-pixel width line scan. **B.** Averaged intensities of the line scans shown in **A.** The intensity of each channel is shown as a differently coloured line: Cell Mask in red, ANAP in blue and GFP in green. The notches on the x-axis mark the location of the plasma membrane. **C.** Two separate western blots against Kir6.2*-HA (left) and Kir6.2*-HA-GFP (right) constructs. Cells were co-transfected with pANAP, eRF1-E55D, and SUR1 unless otherwise indicated. Full-length Kir6.2 constructs are indicated on each gel with a dashed box. **D.** Each lane from the Kir6.2*-HA-GFP gel is displayed normalised to its highest intensity accompanied by the line averaged density trace. The density peak corresponding to ANAP-labelled Kir6.2 is filled in blue. The density peak for C-terminally truncated Kir6.2 is filled in orange. **E.** Chemiluminescence-based surface expression assay for Kir6.2-HA constructs. Each data point represents an individual coverslip of transfected HEK-293T cells, n = 3-6 for each condition. Note the logarithmic scale on the vertical axis.

**Figure 1 - figure supplement 2.**
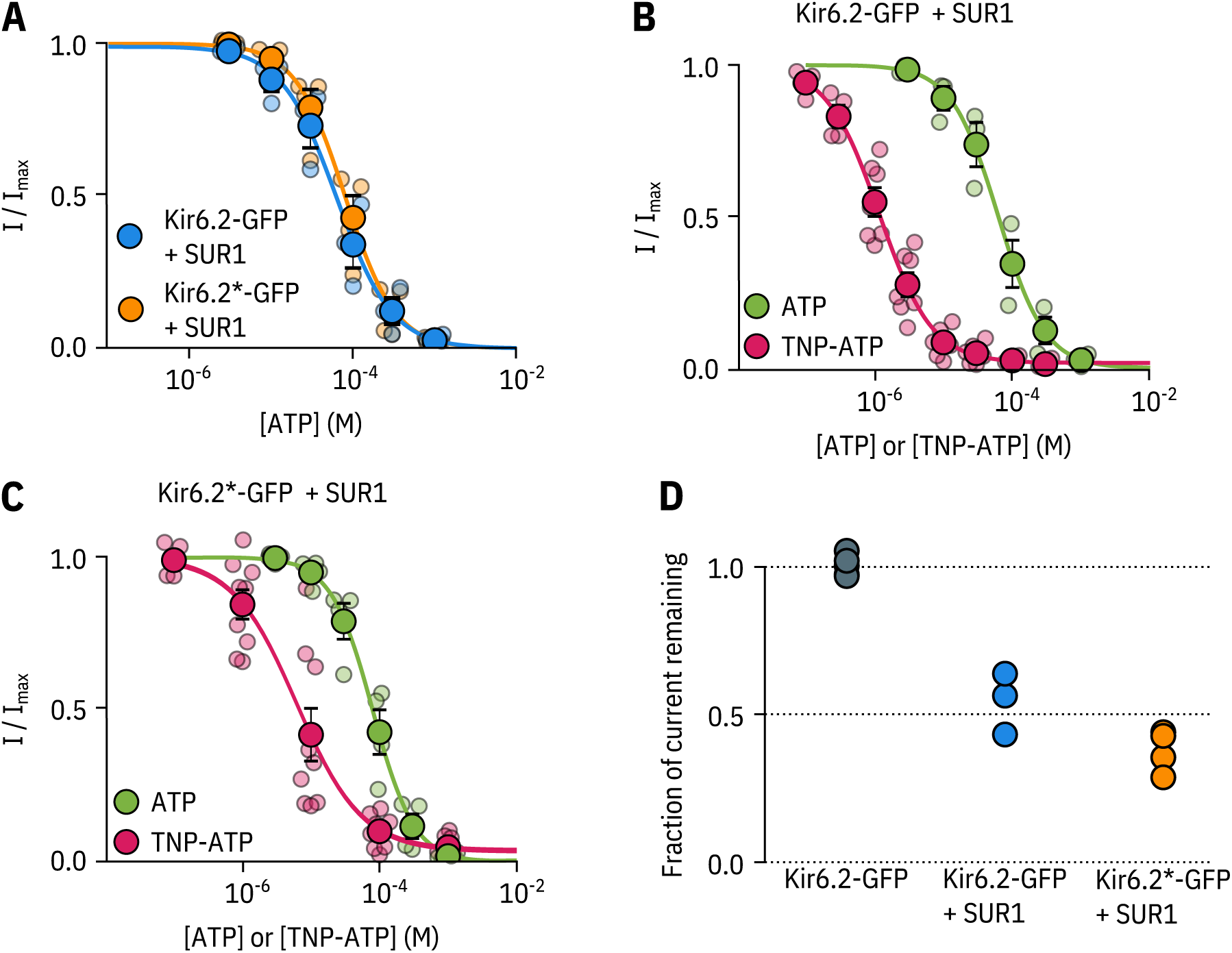
Kir6.2*-GFP is functionally similar to Kir6.2-GFP. **A.** Concentration-response curve for ATP inhibition of Kir6.2-GFP + SUR1 or Kir6.2*-GFP + SUR1, measured in excised, inside-out patches. The smooth curves are descriptive Hill fits to the data, Kir6.2-GFP + SUR1: *IC*_50_ *=* 62.7 μΜ, *h =* 1.28, *I_max_* = 0.99, n = 3; Kir6,2*-GFP + SUR1: *IC*_50_ *=* 79.5 μΜ, *h =* 1.42, *I_max_* = 1.00, n = 4. **B, C.** Concentration-response relationships for current inhibition in excised, inside-out patches expressing Kir6.2-GFP + SUR1 **(C)** or Kir6.2*-GFP + SUR1 (D) exposed to either ATP or TNP-ATP. The smooth curves are descriptive Hill fits to the data, Kir6.2-GFP + SUR1 (TNP-ATP): *IC*_50_ = 1,17μΜ, *h =* 1.14, *I_max_* = 0.97, n = 7, Kir6.2*-GFP + SUR1 (TNP-ATP): *IC*_50_ = 6.23 μΜ, *h =* 0.92, *I_max_ =* 0.96, n = 9. Data and fits for inhibition of Kir6.2*-GFP + SUR1 by TNP-ATP are the same as in Figure 2. **D.** Fractional current inhibition by 100μΜ tolbutamide measured in excised, inside out patches. Data were normalised to the average current in control solution before and after tolbutamide exposure. Each data point represents an individual patch. Kir6.2-GFP without SUR1, n = 5; Kir6.2-GFP + SUR1, n = 3; Kir6,2*-GFP + SUR1, n =4.

**Figure 2 - figure supplement 1.**
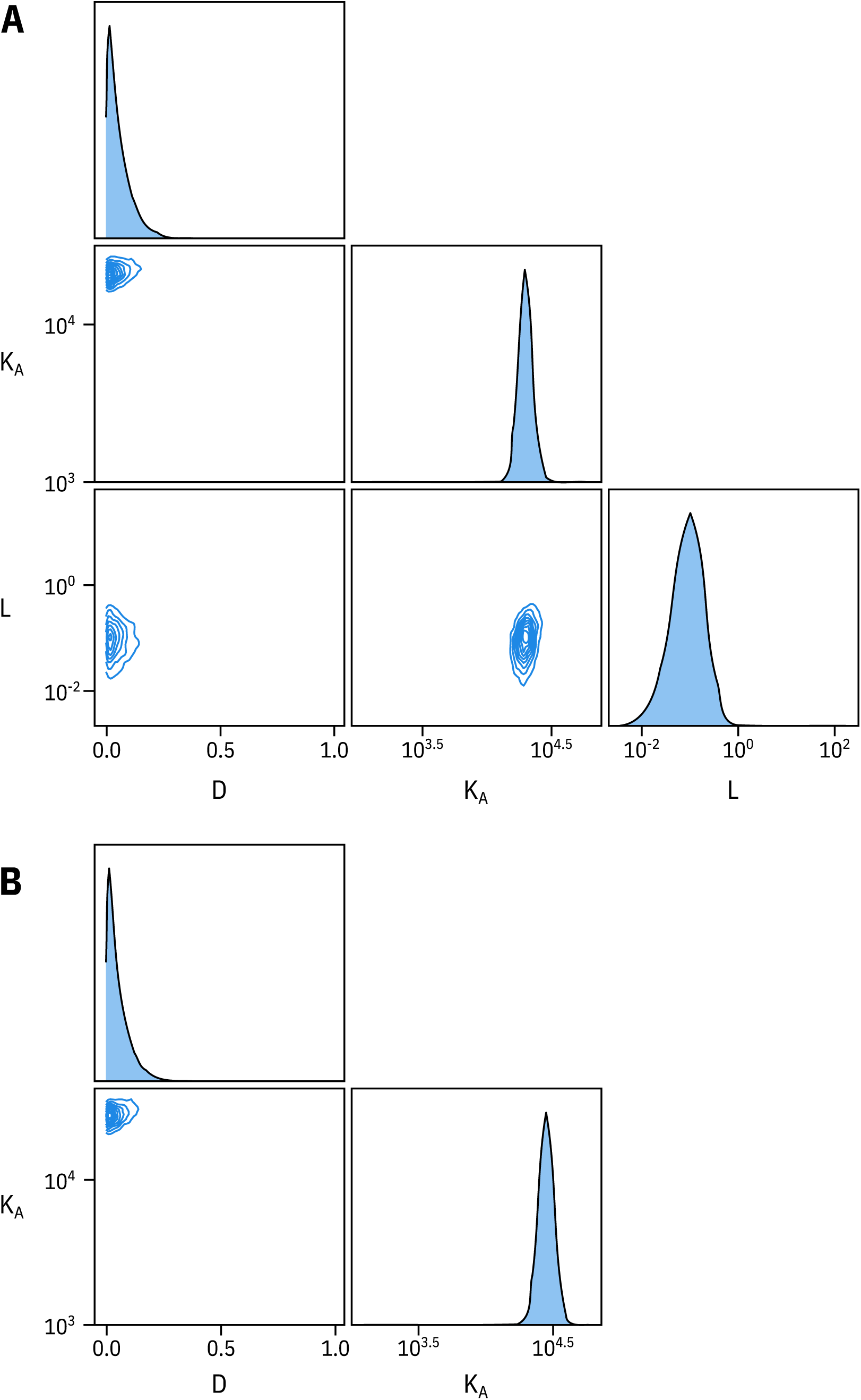
Fixing *L* does not affect estimates of *D* and *K_A_*. **A.** Pairwise correlation plots of *L, D* and *K_A_* from the full MWC-type model fit to Kir6.2*-GFP + SUR1. **B.** Pairwise correlation plots of *D* and *K_A_* from the full MWC-type with *L* fixed to 0.8 (*P_open_ =* 0.45).

**Figure 2 - figure supplement 2.**
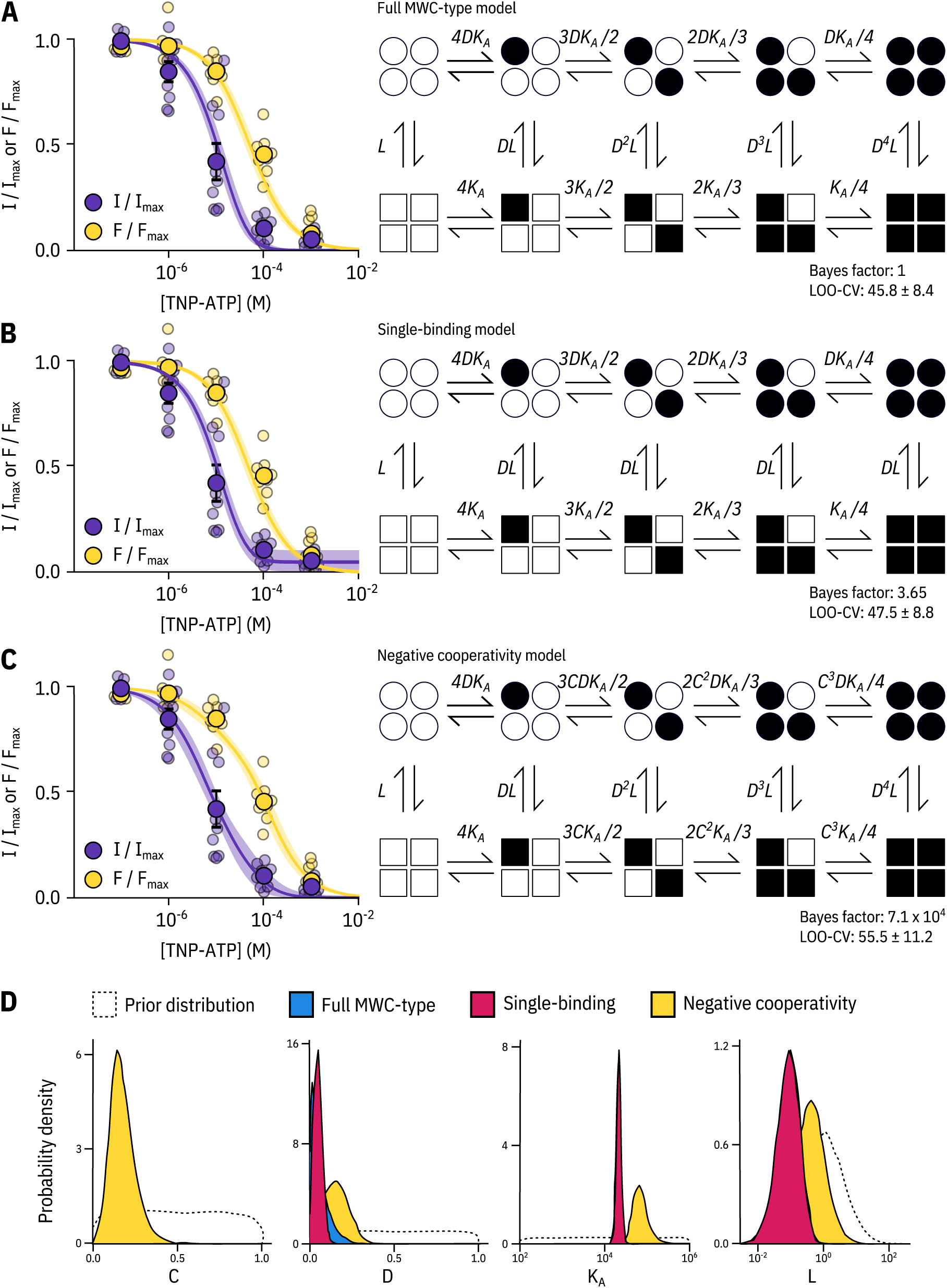
Model selection. Fits to PCF data from Figure 2 with the full MWC-type model **(A),** single-binding model **(B)** and negative-cooperativity model **(C)** are shown on the left with the diagrammatic formulation of each model on the right. The Bayes factor and leave-one-out cross-validation (LOO-CV) scores for each model compared to the full MWC-type model are displayed. *L, D, C*, and *K_A_* are defined in the text. **D.** Posterior probability distributions for the each of the models generated by MCMC fits to the data in Figure 2 overlaid on the prior probability distribution (dashed line) for each parameter. For *L* and *K_A_*, the distributions for the MWC-type and single-binding model were virtually identical. The MWC-type densities are hidden behind the single-binding densities.

**Figure 3 - figure supplement 1.**
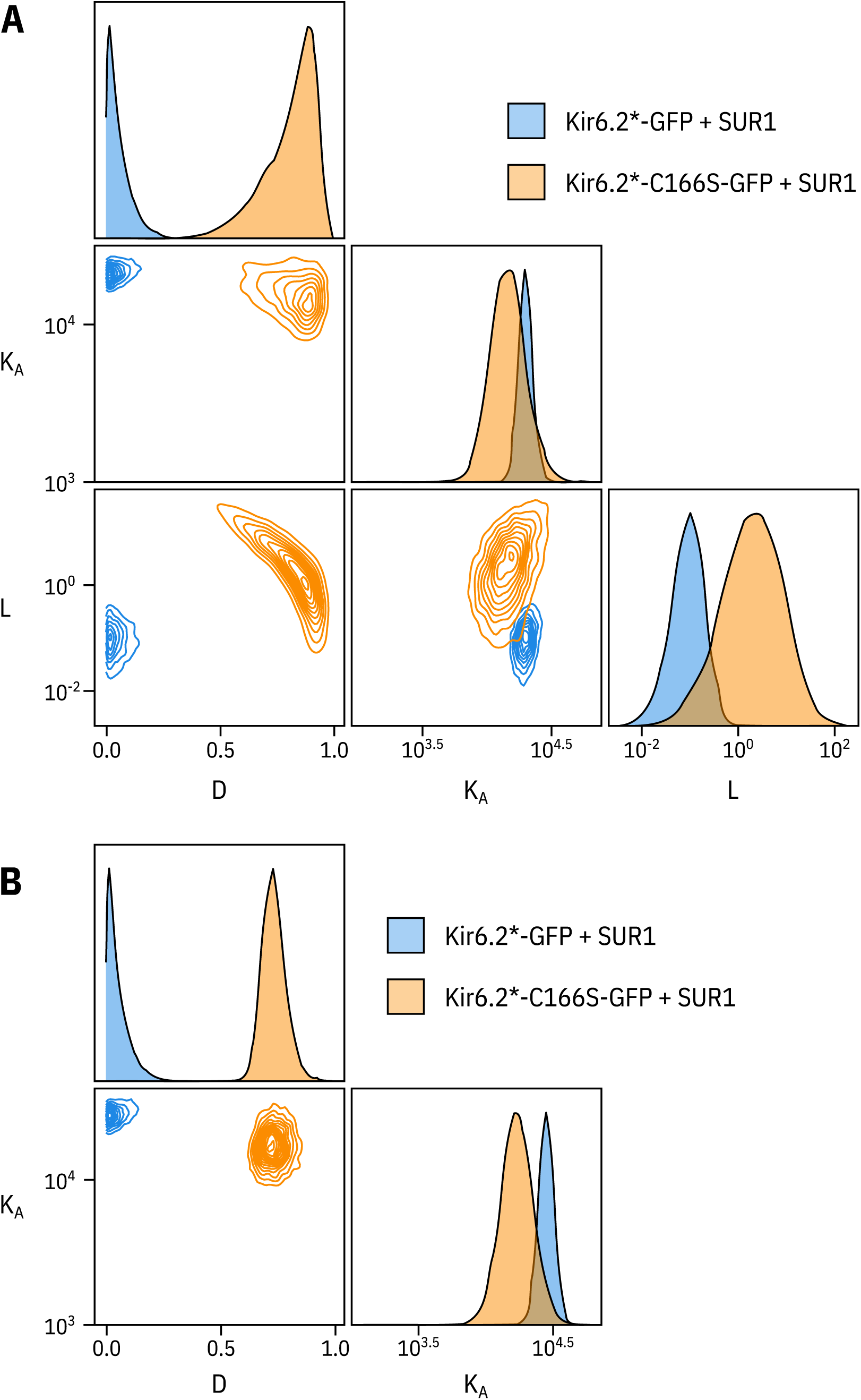
Fixing *L* does not affect the other two parameters. **A.** Pairwise correlation plots of *L, D* and *K_A_* from the full MWC-type model fit to Kir6.2*-GFP + SUR1 and Kir6.2*-C166S-GFP + SUR1. **B.** Pairwise correlation plots of *D* and *K_A_* from the full MWC-type with *L* fixed to 0.8 for Kir6,2*-GFP + SUR1 (*P_open_* = 0.45) or 6.0 for Kir6.2*-C166S-GFP + SUR1 (*P_open_* = 0.86).

**Figure 5 - figure supplement 1.**
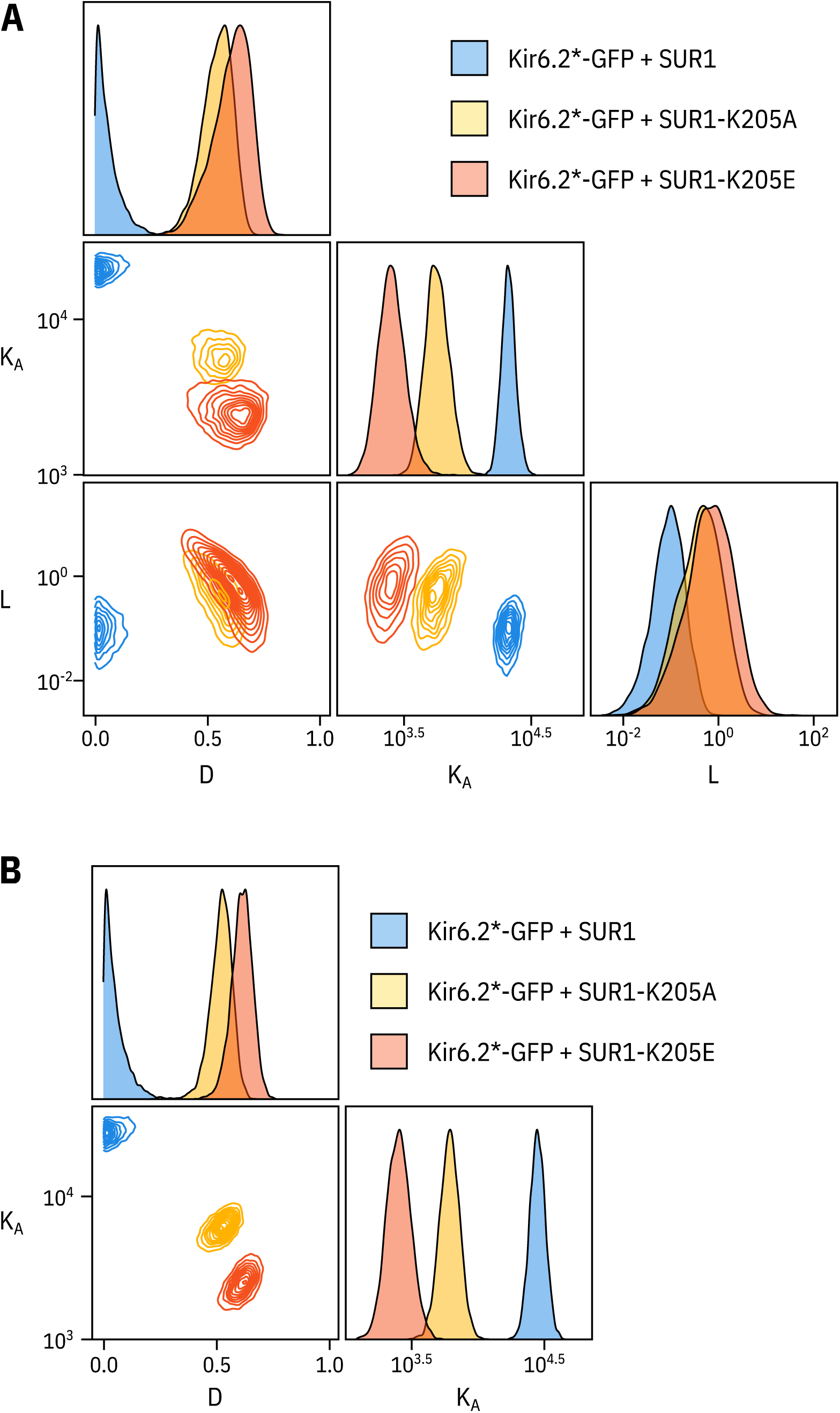
Fixing the *L* parameter does not drastically affect the fits to the SUR1-K205A or SUR1-K205E data. **A.** Pairwise correlation plots of *L, D* and *K_A_* from the full MWC-type model fit to Kir6.2*-GFP co-expressed with wild-type SUR1, SUR1-K205A, and SUR1-K205E. **B.** Pairwise correlation plots of P and *K_A_* from the full MWC-type as above with *L* fixed to 0,8 **(Trapp et al., 1998).**

**Figure 5 - figure supplement 2.**
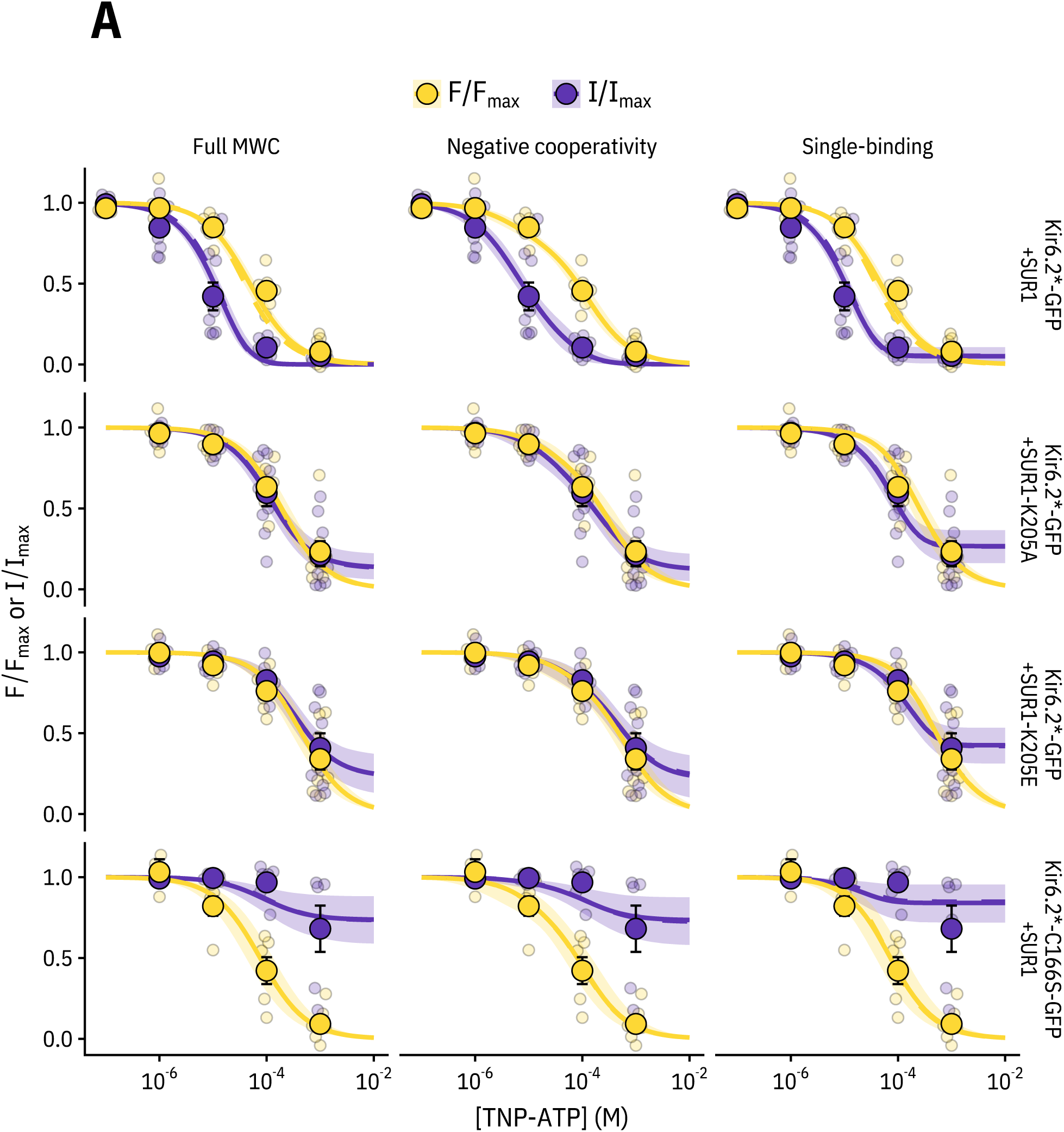
Comparing the ability of each model to explain the data. Fits for each construct with each model (MWC-type, single-binding, negative-cooperativity) are displayed with the solid curve representing the median fit, the shaded area representing the 95% quartiles, and the dashed curve representing the median fit if the *L* parameter is fixed (to 6.0 for Kir6.2*-C166S-GFP + SUR1 and to 0.8 for the other three constructs). As the two fits were very similar, the dashed curve mostly overlays the solid curve. The most notable differences between the fits are that the negative cooperativity model allows for non-sigmoidal curves, and the single-binding model predicts much larger pedestals of current at saturating concentrations of TNP-ATP than either of the other two models.

## Notes

#### Summary of Updates

Minor text revisions.

## References

Aguilar-Bryan L, Nichols CG, Wechsler SW, Clement JPt, Boyd r A E, Gonzalez G, Herrera-Sosa H, Nguy K, Bryan J, Nelson DA. Cloning of the beta ceil high-affinity sulfonylurea receptor: a regulator of insulin secretion. Science. 1995; 268(5209):423–6. https://www.ncbi.nlm.nih.gov/pubmed/7716547.

Ashcroft FM, Puljung MC, Vedovato N. Neonatal Diabetes and the KATP Channel: From Mutation to Therapy. Trends Endocrinol Metab. 2017; 28(5):377–387. https://www.ncbi.nlm.nih.gov/pubmed/28262438, doi: 10.1016/j.tem.2017.02.003.

Ashcroft FM, Rorsman P. K(ATP) channels and islet hormone secretion: new insights and controversies. Nat Rev Endocrinol. 2013; 9(11):660–9. https://www.ncbi.nlm.nih.gov/pubmed/24042324, doi: 10.1038/nrendo.2013.166.

Babenko AP, Bryan j. Sur domains that associate with and gate KATP pores define a novel gatekeeper. J Biol Chem. 2003; 278(43):41577–80. https://www.ncbi.nlm.nih.gov/pubmed/12941953, doi: 10.1074/jbc.C300363200.

Biskup C, Kusch J, Schulz E, Nache V, Schwede F, Lehmann F, Hagen V, Benndorf K. Relating ligand binding to activation gating in CNGA2 channels. Nature. 2007; 446(7134):440–3, https://www.ncbi.nlm.nih.gov/pubmed/17322905, doi: 10.1038/nature05596.

Burkner PC. brms: An R Package for Bayesian Multilevel Models Using Stan. Journal of Statistical Software. 2017; 80(1):1–28. <GotoISI> ://W0S =000408409900001.

Chan KW, Zhang H, Logothetis DE. N-terminal transmembrane domain of the SUR controls trafficking and gating of Kir6 channel subunits. EMBOJ. 2003; 22(15):3833–43. https://www.ncbi.nlm.nih.gov/pubmed/12881418, doi: 10.1093/emboj/cdg376.

Chatterjee A, Guo J, Lee HS, Schultz PG. A genetically encoded fluorescent probe in mammalian cells. J Am Chem Soc. 2013; 135(34):12540–3. https://www.ncbi.nlm.nih.gov/pubmed/23924161, doi: 10.1021/ja4059553.

Craig TJ, Ashcroft FM, Proks P. How ATP inhibits the open K(ATP) channel. J Gen Physiol. 2008; 132(1): 131–44. https://www.ncbi.nlm.NIH.gov/pubmed/18591420, doi: 10.1085/jgp.200709874.

Dolinsky TJ, Nielsen JE, McCammon JA, Baker NA. PDB2PQR: an automated pipeline for the setup of Poisson-Boltzmann electrostatics calculations. Nucleic Acids Res. 2004; 32(Web Server issue):W665–7. https://www.ncbi.nlm.nih.gov/pubmed/15215472, doi: 10.1093/nar/gkh381.

Drain P, Geng X, Li L. Concerted gating mechanism underlying KATP channel inhibition by ATP. BiophysJ. 2004; 86(4):2101–12. https://www.ncbi.nlm.nih.gov/pubmed/15041650, doi: 10.1016/S0006-3495(04)74269-1.

Drain P, Li L, Wang J. KATP channel inhibition by ATP requires distinct functional domains of the cytoplasmic C terminus of the pore-forming subunit. Proc Natl Acad Sci USA. 1998; 95(23): 13953–8. https://www.ncbi.nlm.nih.gov/pubmed/9811907.

Enkvetchakul D, Loussouarn G, Makhina E, Shyng SL, Nichols CG. The kinetic and physical basis of K(ATP) channel gating: toward a unified molecular understanding. BiophysJ. 2000; 78(5):2334–48. https://www.ncbi.nlm.nih.gov/pubmed/10777731, doi: 10.1016/S0006-3495(00)76779-8.

Enkvetchakul D, Nichols CG. Gating mechanism of KATP channels: function fits form. J Gen Physiol. 2003; 122(5):471–80. https://www.ncbi.nlm.nih.gov/pubmed/14581579, doi: 10.1085/jgp.200308878.

Fang K, Csanady L, Chan KW. The N-terminal transmembrane domain (TMDO) and a cytosolic linker (LO) of sulphonyiurea receptor define the unique intrinsic gating of KATP channels. J Physiol. 2006; 576(Pt 2):379–89. https://www.ncbi.nlm.nih.gov/pubmed/16887879, doi: 10.1113/jphysiol.2006.112748.

Flanagan SE, Edghill EL, Gloyn AL, Ellard S, Hattersley AT. Mutations in KCNJ11, which encodes Kir6.2, are a common cause of diabetes diagnosed in the first 6 months of life, with the phenotype determined by genotype. Di-abetologia. 2006; 49(6):1190–7. https://www.ncbi.nlm.nih.gov/pubmed/16609879, doi: 10.1007/s00125-006-0246-z.

Gelman A, Lee D, Guo JQ. Stan: A Probabilistic Programming Language for Bayesian Inference and Optimization. journal of Educational and Behavioral Statistics. 2015; 40(5):530–543. <GotoISI> ://W0S: 000363883900006.

Gloyn AL, Diatloff-Zito C, Edghill EL, Bellanne-Chantelot C, Nivot S, Coûtant R, Ellard S, Hattersley AT, Robert JJ. KCNJ11 activating mutations are associated with developmental delay, epilepsy and neonatal diabetes syndrome and other neurological features. EurJ Hum Genet. 2006; 14(7):824–30. https://www.ncbi.nlm.nih.gov/pubmed/16670688, doi: 10.1038/sj.ejhg.5201629.

Gribble FM, Tucker SJ, Haug T, Ashcroft FM, MgATP activates the beta cell KATP channel by interaction with its SUR1 subunit, Proc Natl Acad Sci USA, 1998; 95(12):7185–90. https://www.ncbi.nlm.nih.gov/pubmed/9618560, doi: 10.1073/pnas.95.12.7185.

Gronau QF, Singmann H, Wagenmakers EJ. bridgesampling; An R Package for Estimating Normalizing Constants, arXiv e-prints. 2017 Oct; p. arXiv:1710.08162.

Heuser J. The production of’cell cortices’for light and electron microscopy. Traffic. 2000; 1(7):545–52. https://www.ncbi.nlm.nih.gov/pubmed/11208142.

Hines KE, Middendorf TR, Aldrich RW. Determination of parameter identifiability in nonlinear biophysical models: A Bayesian approach. J Gen Physiol. 2014; 143(3):401–16. https://www.ncbi.nlm.nih.gov/pubmed/24516188, doi: 10.1085/jgp.201311116.

Inagaki N, Gonoi T, Clement JPt, Namba N, Inazawa J, Gonzalez G, Aguilar-Bryan L, Seino S, Bryan J. Reconstitution of I KATP: an inward rectifier subunit plus the sulfonylurea receptor. Science. 1995; 270(5239): 1166–70, https://www.ncbi.nlm.nih.gov/pubmed/7502040.

Inagaki N, Gonoi T, Seino S. Subunit stoichiometry of the pancreatic beta-cell ATP-sensitive K+ channel. FEBS Lett. 1997; 409(2)232–6. https://www.ncbi.nlm.nih.gov/pubmed/9202152.

John SA, Monck JR, Weiss JN, Ribalet B. The sulphonylurea receptor SUR1 regulates ATP-sensitive mouse Kir6.2 K+ channels linked to the green fluorescent protein in human embryonic kidney cells (HEK 293). J Physiol. 1998; 510 (Pt2):333–45. https://www.ncbi.nlm.nih.gov/pubmed/9705987, doi: 10.1111/j.1469-7793.1998.333bk.x.

Kasuya G, Yamaura T, Ma XB, Nakamura R, Takemoto M, Nagumo H, Tanaka E, Dohmae N, Nakane T, Yu Y, Ishitani R, Matsuzaki O, Hattori M, Nureki O. Structural insights into the competitive inhibition of the ATP-gated P2X receptor channel. Nat Commun. 2017; 8(1):876. https://www.ncbi.nlm.nih.gov/pubmed/29026074, doi: 10.1038/S41467-017-00887-9.

Kusch J, Biskup C, Thon S, Schulz E, Nache V, Zimmer T, Schwede F, Benndorf K. Interdependence of receptor activation and ligand binding in HCN2 pacemaker channels. Neuron. 2010; 67(1):75–85. https://www.ncbi.nlm.nih.gov/pubmed/20624593, doi: 10.1016/j.neuron.2010.05.022.

Li L, Wang J, Drain P. The 1182 region of k(ir)6.2 is closely associated with ligand binding in K(ATP) channel inhibition by ATP. Biophys J. 2000; 79(2):841–52. https://www.ncbi.nlm.nih.gov/pubmed/10920016, doi: 10.1016/S0006-3495(00)76340-5.

Lin CY, Huang Z, Wen W, Wu A, Wang C, Niu L. Enhancing Protein Expression in HEK-293 Cells by Lowering Culture Temperature. PLoS One. 2015; 10(4):e0123562. https://www.ncbi.nlm.nih.gov/pubmed/25893827, doi: 10.1371/journal.pone.0123562.

Makhina EN, Nichols CG. Independent trafficking of KATP channel subunits to the plasma membrane. J Biol Chem. 1998; 273(6):3369–74. https://www.ncbi.nlm.nih.gov/pubmed/9452456.

Markworth E, Schwanstecher C, Schwanstecher M, ATP4-mediates closure of pancreatic beta-cell ATP-sensitive potassium channels by interaction with 1 of 4 identical sites. Diabetes. 2000; 49(9): 1413–8. https://www.ncbi.nlm.nih.gov/pubmed/10969823, doi: 10.2337/diabetes.49.9.1413.

Martin GM, Kandasamy B, DiMaio F, Yoshioka C, Shyng SL. Anti-diabetic drug binding site in a mammalian KATP channel revealed byCryo-EM. Elife. 2017; 6. https://www.ncbi.nlm.nih.gov/pubmed/29035201, doi: 10.7554/eLife.31054.

Martin GM, Sung MW, Yang Z, Innés LM, Kandasamy B, David LL, Yoshioka C, Shyng SL. Mechanism of pharma-cochaperoning in a mammalian KATP channel revealed by cryo-EM. Elife. 2019; 8. https://www.ncbi.nlm.nih.gov/pubmed/31343405, doi: 10.7554/eLife.46417.

Masia R, Koster J C, Tumini S, Chiarelli F, Colombo C, Nichols CG, Barbetti F. An ATP-binding mutation (G334D) in KCNJ11 is associated with a sulfonylurea-insensitive form of developmental delay, epilepsy, and neonatal diabetes. Diabetes. 2007; 56(2):328–36. https://www.ncbi.nlm.nih.gov/pubmed/17259376, doi: 10.2337/db06-1275.

McTaggart JS, Clark RH, Ashcroft FM. The role of the KATP channel in glucose homeostasis in health and disease: more than meets the islet, J Physiol. 2010; 588(Pt 17):3201–9. https://www.ncbi.nlm.nih.gov/pubmed/20519313, doi: 10.1113/jphysiol.2010.191767.

Monod J, Wyman J, Changeux JP. On the Nature of Allosteric Transitions: A Plausible Model. J Mol Biol. 1965; 12:88–118. https://www.ncbi.nlm.nih.gov/pubmed/14343300.

Nichols CG, Shyng SL, Nestorowicz A, Glaser B, Clement JPt, Gonzalez G, Aguilar-Bryan L, Permutt MA, Bryan J. Adenosine diphosphate as an intracellular regulator of insulin secretion. Science. 1996; 272(5269): 1785–7. https://www.ncbi.nlm.nih.gov/pubmed/8650576.

Pratt EB, Zhou Q, Gay JW, Shyng SL. Engineered interaction between SUR1 and Kir6.2 that enhances ATP sensitivity in KATP channels, J Gen Physiol. 2012; 140(2):175–87. https://www.ncbi.nlm.nih.gov/pubmed/22802363, doi: 10.1085/jgp.201210803.

Proks P, Puljung MC, Vedovato N, Sachse G, Mulvaney R, Ashcroft FM. Running out of time: the decline of channel activity and nucleotide activation in adenosine triphosphate-sensitive K-channels. Philos Trans R Soc Lond B Biol Sci. 2016; 371(1700). https://www.ncbi.nlm.nih.gov/pubmed/27377720, doi: 10.1098/rstb.2015.0426.

Proks P, de Wet H, Ashcroft FM. Activation of the K(ATP) channel by Mg-nucleotide interaction with SUR1. J Gen Physiol. 2010; 136(4):389–405. https://www.ncbi.nlm.nih.gov/pubmed/20876358, doi: 10.1085/jgp.201010475.

Puljung M, Vedovato N, Usher S, Ashcroft F. Activation mechanism of ATP-sensitive K(+) channels explored with real-time nucleotide binding. Elife. 2019; 8. https://www.ncbi.nlm.nih.gov/pubmed/30789344, doi: 10.7554/eLife.41103.

Puljung MC. Cryo-electron microscopy structures and progress toward a dynamic understanding of KATP channels. J Gen Physiol. 2018; 150(5):653–669. https://www.ncbi.nlm.nih.gov/pubmed/29685928, doi: 10.1085/jgp.201711978.

Quan Y, Barszczyk A, Feng ZP, Sun HS. Current understanding of KATP channels in neonatal diseases: focus on insulin secretion disorders. Acta Pharmacol Sin. 2011; 32(6):765–80. https://www.ncbi.nlm.nih.gov/pubmed/21602835, doi; 10.1038/aps.2011.57.

Ribalet B, John SA, Xie LH, Weiss JN. ATP-sensitive K+ channels: regulation of bursting by the sulphonylurea receptor, PIP2 and regions of Kir6.2. J Physiol. 2006; 571 (Pt 2):303–17. https://www.ncbi.nlm.nih.gov/pubmed/16373383, doi: 10.1113/jphysiol.2005.100719.

Sakura H, Ammala C, Smith PA, Gribble FM, Ashcroft FM. Cloning and functional expression of the cDNA encoding a novel ATP-sensitive potassium channel subunit expressed in pancreatic beta-cells, brain, heart and skeletal muscle. FEBS Lett. 1995; 377(3):338–44. https://www.ncbi.nlm.nih.gov/pubmed/8549751, doi: 10.1016/0014-5793(95)01369-5.

Schmied WH, Elsasser SJ, Uttamapinant C, Chin JW. Efficient muitisite unnatural amino acid incorporation in mammalian cells via optimized pyrrolysyl tRNA synthetase/tRNA expression and engineered eRF1. J Am Chem Soc. 2014; 136(44):15577–83. https://www.ncbi.nlm.nih.gov/pubmed/25350841, doi: 10.1021/ja5069728.

Snider KE, Becker S, Boyajian L, Shyng SL, MacMullen C, Hughes N, Ganapathy K, Bhatti T, Stanley CA, Ganguly A. Genotype and phenotype correlations in 417 children with congenital hyperinsulinism. J Clin Endocrinol Metab. 2013; 98(2):E355–63. https://www.ncbi.nlm.nih.gov/pubmed/23275527, doi: 10.1210/jc.2012-2169.

Toyoshima C, Yonekura S, Tsueda J, Iwasawa S. Trinitrophenyl derivatives bind differently from parent adenine nucleotides to Ca2+-ATPase in the absence of Ca2+. Proc Natl Acad Sci USA. 2011; 108(5):1833–8. https://www.ncbi.nlm.nih.gov/pubmed/21239683, doi: 10.1073/pnas.1017659108.

Trapp S, Proks P, Tucker SJ, Ashcroft FM. Molecular analysis of ATP-sensitive K channel gating and implications forchannel inhibition by ATP. J Gen Physiol. 1998; 112(3):333–49. https://www.ncbi.nlm.nih.gov/pubmed/9725893, doi: 10.1085/jgp.112.3.333.

Trott O, Olson AJ, AutoDock Vina: improving the speed and accuracy of docking with a new scoring function, efficient optimization, and multithreading. J Comput Chem. 2010 Jan; 31 (2):455–461.

Tucker SJ, Gribble FM, Zhao C, Trapp S, Ashcroft FM. Truncation of Kir6.2 produces ATP-sensitive K+ channels in the absence of the sulphonylurea receptor. Nature. 1997; 387(6629): 179–83. https://www.ncbi.nlm.nih.gov/pubmed/9144288, doi: 10.1038/387179a0.

Tusnady GE, Bakos E, Varadi A, Sarkadi B. Membrane topology distinguishes a subfamily of the ATP-binding cassette (ABC) transporters. FEBS Lett. 1997; 402(1):1–3. https://www.ncbi.nlm.nih.gov/pubmed/9013845, doi: 10.1016/s0014-5793(96)01478-0.

Ueda K, Komine J, Matsuo M, Seino S, Amachi T. Cooperative binding of ATP and MgADP in the sulfonylurea receptor is modulated byglibenclamide. Proc Natl Acad Sci USA. 1999; 96(4): 1268–72. https://www.ncbi.nlm.nih.gov/pubmed/9990013.

Vedovato N, Ashcroft FM, Puljung MC. The Nucleotide-Binding Sites of SUR1: A Mechanistic Model. Biophys J. 2015; 109(12):2452–60. https://www.ncbi.nlm.nih.gov/pubmed/26682803, doi: 10.1016/j.bpj.2O15.10.026.

Vehtari A, Gelman A, Gabry J. Practical Bayesian model evaluation using leave-one-out cross-validation and WAIC (vol 27, pg 1413, 2017). Statistics and Computing. 2017; 27(5): 1433–1433. <GotoISI>://WOS: 000400831700018, doi: 10.1007/s11222-016-9709-3.

Wagenmakers EJ. A practical solution to the pervasive problemsofp values. Psychon Bull Rev. 2007; 14(5):779–804. https://www.ncbi.nlm.nih.gov/pubmed/18087943.

Wang R, Zhang X, Cui N, Wu J, Piao H, Wang X, Su J, Jiang C. Subunit-stoichiometric evidence for kir6.2 channel gating, ATP binding, and binding-gating coupling. Mol Pharmacol. 2007; 71 (6): 1646–56. https://www.ncbi.nlm.nih.gov/pubmed/17369308, doi: 10.1124/mo1.106.030528.

Wu S, Vysotskaya ZV, Xu X, Xie C, Liu Q, Zhou L. State-dependent cAMP binding to functioning HCN channels studied by patch-ciamp fluorometry. Biophys j. 2011; 100(5): 1226–32. https://www.ncbi.nlm.nih.gov/pubmed/21354395, doi: 10.1016/j.bpj.2011.01.034.

Yan FF, Lin YW, MacMullen C, Ganguly A, Stanley CA, Shyng SL. Congenital hyperinsulinism associated ABCC8 mutations that cause defective trafficking of ATP-sensitive K+channels: identification and rescue. Diabetes. 2007; 56(9):2339–48. https://www.ncbi.nlm.nih.gov/pubmed/17575084, doi: 10.2337/db07-0150.

Zagotta WN, Gordon MT, Senning EN, Munari MA, Gordon SE. Measuring distances between TRPV1 and the plasma membrane using a noncanonical amino acid and transition metal ion FRET, J Gen Physiol. 2016; 147(2):201–16. https://www.ncbi.nlm.nih.gov/pubmed/26755770, doi: 10.1085/jgp.201511531.

Zerangue N, Schwappach B, Jan YN, Jan LY. A new ER trafficking signal regulates the subunit stoichiometry of plasma membrane K(ATP) channels. Neuron. 1999; 22(3):537–48. https://www.ncbi.nlm.nih.gov/pubmed/10197533.

Zheng J, Zagotta WN. Patch-clamp fluorometry recording of conformational rearrangements of ion channels. Sci STKE. 2003; 2003(176):PL7. https://www.ncbi.nlm.nih.gov/pubmed/12671191, doi: 10.1126/stke.2003.176.pl7.

